# A Genetic Model Therapy Proposes a Critical Role for Liver Dysfunction in Mitochondrial Biology and Disease

**DOI:** 10.1101/2020.05.08.084681

**Authors:** Ankit Sabharwal, Mark D. Wishman, Roberto Lopez Cervera, MaKayla R. Serres, Jennifer L. Anderson, Anthony J. Treichel, Noriko Ichino, Weibin Liu, Jingchun Yang, Yonghe Ding, Yun Deng, Steven A. Farber, Karl J. Clark, Xiaolei Xu, Stephen C. Ekker

## Abstract

The clinical and largely unpredictable heterogeneity of phenotypes in patients with mitochondrial disorders demonstrates the ongoing challenges in the understanding of this semi-autonomous organelle in biology and disease. Here we present a new animal model that recapitulates key components of Leigh Syndrome, French Canadian Type (LSFC), a mitochondrial disorder that includes diagnostic liver dysfunction. LSFC is caused by allelic variations in the Leucine Rich Pentatricopeptide repeat-containing motif (*LRPPRC*) gene. *LRPPRC* has native functions related to mitochondrial mRNA polyadenylation and translation as well as a role in gluconeogenesis. We used the Gene-Breaking Transposon (GBT) cassette to create a revertible, insertional mutant zebrafish line in the *LRPPRC* gene. *lrpprc* zebrafish homozygous mutants displayed impaired muscle development, liver function and lowered levels of mtDNA transcripts and are lethal by 12dpf, all outcomes similar to clinical phenotypes observed in patients. Investigations using an *in vivo* lipidomics approach demonstrated accumulation of non-polar lipids in these animals. Transcript profiling of the mutants revealed dysregulation of clinically important nuclearly encoded and mitochondrial transcripts. Using engineered liver-specific rescue as a genetic model therapy, we demonstrate survival past the initial larval lethality, as well as restored normal gut development, mitochondrial morphology and triglyceride levels functionally demonstrating a critical role for the liver in the pathophysiology of this model of mitochondrial disease. Understanding the molecular mechanism of the liver-mediated genetic rescue underscores the potential to improve the clinical diagnostic and therapeutic developments for patients suffering from these devastating disorders.

## Introduction

The mitochondrion is a complex and essential organelle whose dysfunction is linked to a panoply of diverse human pathologies and diseases. The understanding of the traditional role of mitochondria as the powerhouses of the cell has evolved as the many roles in cellular and physiological homeostasis have been functionally enumerated far beyond this commonly known role in energy production via the electron transport chain (ETC). Today, mitochondria are known to directly contribute to calcium signaling, apoptosis, iron homeostasis, lipid metabolism, ATP-metabolism, immunity, and a host of other critical biochemical synthesis pathways (Green, 1998; Rizzuto et al., 2009; Spinelli and Haigis, 2018; Tiku et al., 2020; Valero, 2014). All of these cellular functions rely on well-orchestrated cross-talk between the nuclear and mitochondrial genome. Genetic lesions arising in the nuclear and/or mitochondrial genome can lead to mitochondrial dysfunction that present in humans with heterogeneous groups of clinical manifestations that differentially impact every organ system in the body (Chinnery et al., 2012; Vafai and Mootha, 2012). The widespread activity of mitochondria provides ample opportunities for mitochondrial dysfunction to play a role in human disease, but current understanding does not provide specificity in differential biological function or capacity in accurately predicting pathogenic details suitable for therapeutic development.

Mitochondrial disease can arise directly such as pathogenic genetic variations in mitochondrial DNA such as Mitochondrial Encephalopathy Lactic acidosis and Stroke (MELAS) or nuclear DNA including Leigh Syndrome, or as a secondary condition associated with an underlying pathology such as Alzheimer’s disease, diabetes, or cancer (Moraes et al., 1992; Newsholme et al., 2012; Rahman et al., 1996; Urrutia et al., 2014; Wallace, 2012). Even in these circumstances where the proximal cause of disease initiation is known, the specific manifestations of mitochondrial dysfunction can produce a wide spectrum of clinical features that vary in severity and tissue specificity, even within patients harboring identical genetic variations (Gorman et al., 2016; McFarland et al., 2010). Therefore, establishing causality between phenotypes and genetic variations for a given mitochondrial disease is often challenging in a clinical setting.

Here we focus on Leigh Syndrome, French Canadian Type (LSFC), a well-defined mitochondrial disease with onset in infancy that manifests with diagnostic liver dysfunction. LSFC is a monogenic, autosomal recessive condition. LSFC was first discovered in the Saguenay-Lac-Saint-Jean region of Quebec, Canada, where roughly 1 in 23 individuals was found to be a carrier of a diseased allele (Morin et al., 1993). The most common allelic variant is due to an A354V transition in exon 9 of the leucine rich pentatricopeptide repeat containing motif (*LRPPRC*) protein, a nuclear-encoded gene. Other pathogenic variants in *LRPPRC* have been reported in patients, including an 8 base pair deletion in exon 35 (Mootha et al., 2003) that presents with reduced phenotypes compared to the A354V mutation. Patients with mutations in *LRPPRC* experience a diverse array of clinical features centered around cytochrome oxidase (COX) deficiency. These can include an early onset of hepatic microvesicular steatosis and chronic lactic acidosis as neonates. Individuals born with LSFC have an average lifespan of less than 5 years of age with most succumbing to a series of extreme, acute metabolic crises. Patients that make it past the early life crises show a lessened disease state characterized by hypotonia, language and mobility deficits and delays, and muscle weakness (Debray et al., 2011; Morin et al., 1993). *LRPPRC* has also been implicated in other diseases such as neurofibromatosis, Parkinson’s disease, and viral infections such as HIV-1 (Cui et al., 2019).

LRPPRC belongs to the pentatricopeptide repeat (PPR) containing motif family of proteins. Pentatricopeptide Repeats (PPRs) are non-catalytic RNA binding domains. PPR proteins consist of a series of ∼35 amino acid repeats wherein two hypervariable residues, those in the 5th and 35th position of the repeat, direct the binding of each repeat region to a single specific ribonucleotide (Manna, 2015). The LRPPRC protein in humans has 20 annotated PPR domains and is promiscuous, preferentially binding to different mitochondrial RNAs (Figure 1A). The gene also has tetratricopeptide (TPR)- and HEAT (huntingtin-elongation A subunit-TOR)-like tandem repeat sequences. The N terminus consists of multiple copies of leucine rich nuclear transport signals, four copies of the transcriptional co-regulator signature LXXLL and PPR motifs extending through the C terminus. The C terminus of the protein also consists of ENTH (Epsin1 N-Terminal Homology) involved in cytoskeletal organization, vesicular trafficking, DUF (Domain of Unknown Function) 28 and SEC1 domain for vesicular transport. *LRPPRC* plays roles in the regulation of both nuclear and mitochondrial RNA expression at the transcriptional and post transcriptional level (Liu and McKeehan, 2002; Mili and Piñol-Roma, 2003).

**Figure 1:**
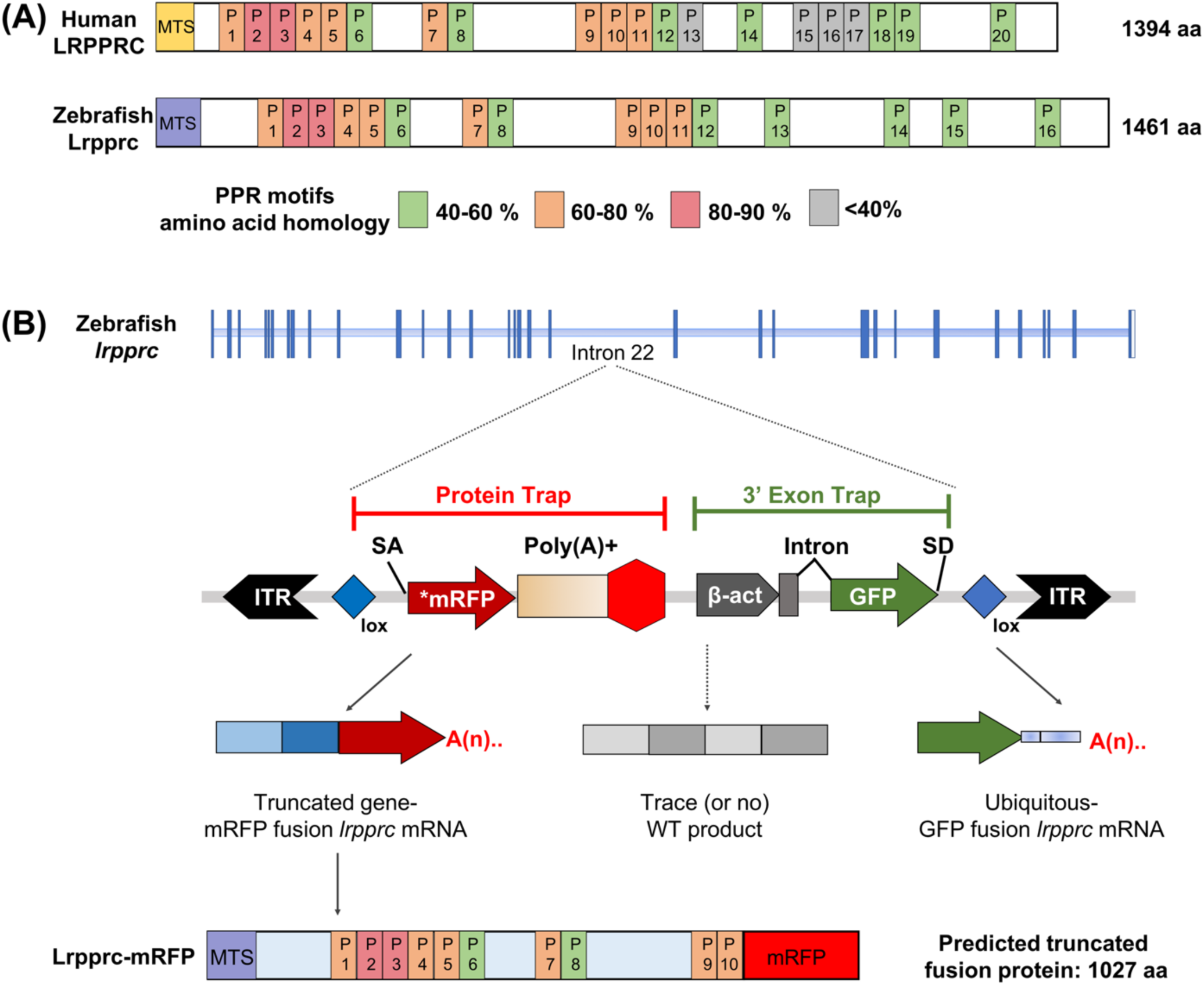
GBT mutagenesis generates a novel zebrafish model of LSFC. **(A)** Schematic of human and zebrafish LRPPRC proteins with highlighted PPR domains (denoted by P). **(B)** Schematic of the integration event of GBT vector RP2.1 with 5’ protein trap and 3’ exon trap cassettes. The RP2.1 cassette was integrated in the intronic region 22 of the lrpprc genomic locus on chromosome 13. ITR, inverted terminal repeat; SA, loxP; Cre recombinase recognition sequence, splice acceptor; *mRFP’ AUG-less mRFP sequence; poly (A)+, polyadenylation signal; red octagon, extra transcriptional terminator and putative border element; β-act, carp beta-actin enhancer, SD, splice donor

The majority of the protein is imported to mitochondria, where it forms a complex with the steroid receptor RNA activator stem-loop (*SLIRP*) protein in the mitochondrial matrix. The absence of *LRPPRC* leads to oligoadenylation of the transcripts. Studies using shRNAs to knockdown *LRPPRC* in MCH58 human fibroblasts showed decreased transcript levels of genes encoded on the mitochondrial chromosome (Gohil et al., 2010). *LRPPRC-SLIRP* is also observed to relax RNA structure, making the 5’ end of the mRNA accessible for the mitochondrial ribosome to initiate translation (Cui et al., 2019). Loss of functional *lrpprc* in *C. elegans* and mice revealed defects in mitochondrial biogenesis, decreased complex 1 and cytochrome c oxidase activity, decreased stability of mitochondrial mRNAs, and dysregulated mitochondrial translation (Cuillerier et al., 2017; Kohler et al., 2015; Xu et al., 2012).

Given that onset of LSFC is marked by the neurological and metabolic crisis (Morin et al., 1993) and the liver is the major metabolic factory of the cell, therefore, the role of liver impairment in LSFC represents a unique aspect of mitochondrial disease unseen in traditional Leigh syndrome. Mice harboring liver-specific inactivation of *Lrpprc* displayed manifestations similar to LSFC patients marked by mitochondrial hepatopathy, growth delay and reduced fatty acid oxidation compared to controls. These animals also exhibited impaired cytochrome c oxidase and ATP synthase activity along with increased susceptibility to calcium-induced permeability transition (Cuillerier et al., 2017). Recently, a lipidomic profile study on LSFC patients highlighted a novel role of this protein in the peroxisomal lipid metabolism (Ruiz et al., 2019).

Current diagnosis of LSFC hinges on measuring lactate levels in the blood and brain, cytochrome C oxidase (COX) activity in patient fibroblasts, and sequencing of the *LRPPRC* gene for mutations (Debray et al., 2011). The lack of a cure or effective therapies for LSFC patients means that current clinicians must shift their focus to relieving and controlling symptoms. Focus on dietary restrictions and lifestyle changes, including exercise and infection risk management, are made to reduce the physiological and metabolic stress on the patient. To build upon a heuristic paradigm for therapeutic interventions for these classes of disorders, we need to establish disease models that serve as a suitable substrate to recapitulate the key functional tissue-specific pathology of the disease.

As a model organism, zebrafish (*Danio rerio*) offers various advantages such as a short breeding cycle, high fecundity, ex utero development, optically clear embryogenesis, rapid development of internal organs, and easy maintenance (Lieschke and Currie, 2007). Zebrafish are a powerful potential vertebrate model organism to study human mitochondrial disorders because of the conserved mitochondrial genome and mitochondrial genetic machinery, as compared to its human counterpart. The zebrafish and human mitochondrial genomes display ∼65% sequence identity at nucleotide level, and share the same codon usage, strand specific nucleotide bias and gene order (Broughton et al., 2001; Sabharwal et al., 2019a). Recently, the zebrafish model has shed light on mitochondrial disorders and helped expand our understanding of the mechanisms of mitochondrial-associated pathology (Byrnes et al., 2018; Flinn et al., 2009; Plucińska et al., 2012; Song et al., 2009; Steele et al., 2014).

We employed an insertional mutagenesis screen using the gene-breaking transposon (GBT) (Clark et al., 2011) to model LSFC. The GBT construct is incorporated into the intron of an endogenous gene via Tol2 transposase and uses a splice acceptor site to create a fusion protein between the native transcript and a monomeric red fluorescent protein (mRFP) gene that contains a transcription termination site (Fig 1B). When inserted intergenically, this result in translation of a truncated endogenous gene product fused to mRFP. This enables spatiotemporal tracking of the trapped gene’s natural protein expression in real time. This insertion is reversible due to two loxP sites located near each terminal repeat that enables removal of the GBT cassette using Cre recombinase. The revertible nature of this construct allows for reversion of the mutant back to wild type alleles to allow for tissue-specific or ubiquitous rescue of the mutated gene (Clark et al., 2011; Ichino et al., 2019).

We describe here the first revertible zebrafish mutant with a single disruptive insertion in the *lrpprc* gene to model LSFC. *lrpprc* homozygous mutants mimic many hallmarks observed in patients such as early lethality, defective muscle development and decreases in mitochondrial transcript levels. In addition, altered dietary lipid metabolism along with mitochondrial dysmorphology was also observed in these mutants. A key outcome of this study is the reversion of the LSFC-associated phenotype and survival using liver-specific Cre recombinase in the homozygous *lrpprc* mutants, demonstrating a critical role of this organ is the pathogenesis of disease in this animal model. This result contributes a novel paradigm towards identification of therapeutic approaches for LSFC and for other related disorders.

## Methods

### Zebrafish handling and husbandry

All adult zebrafish and embryos were maintained according to the guidelines established by Mayo Clinic Institutional Animal Care and Use Committee (Mayo IACUC) (IACUC number: A34513-13-R16).

### Mutant generation

The GBT0235 allele was generated by injecting the pGBT-RP2.1 cassette as described to create the *in vivo* protein trap lines (Clark et al., 2011; Ichino et al., 2019). Generation of these transgenic fish lines involved microinjection of the vector DNA together with the transposase RNA (Tol2) in single cell zebrafish embryos, followed by screening of the injected animals for GFP and RFP expressions. To generate the larvae for liver-specific rescue data, non-GBT transgenic marker lines carrying liver-specific Cre recombinase *Tg(*−*2.8fabp10:Cre;* −*0.8cryaa:Venus)*^*S955*^ were used. *Tg(MLS: EGFP)* transgenic line was used to generate larvae to study the subcellular localization of the truncated fusion protein.

### Zebrafish embryo genotyping (ZEG) protocol

Deficiency of LRPPRC in humans shows an early neonatal phenotype, necessitating the ability to investigate GBT0235 mutants at early stages of life. Likewise, our *lrpprc*^*GBT0235/GBT0235*^ mutants exhibit lethality starting around 8 dpf. We needed the ability to genotype larvae at a young age without sacrificing to enable experiments to be efficiently conducted on living animals such as the survival assays and HPLC lipid analysis. An automated genotyping device was employed for the rapid cellular extraction of DNA from zebrafish embryos (Lambert et al., 2018). Larvae at 3 days post fertilization (dpf) were rinsed three times in fresh embryo water and transferred to a new petri dish. Larvae were aspirated in 12 µL of embryo media and placed on extraction chips in individual wells. An evaporation cover was attached, and the larvae were agitated using a vibrating motor that was powered with 2.4V for 10 minutes. Immediately following the vibration period, the samples were removed from the chip and placed into strip tubes for storage. Larvae were replenished with embryo water and individually transferred to a 24 well plate for storage at 28°C. The collected sample was used as a template for PCR amplification for genotyping of the larvae. Two PCR reactions were run using the same thermal cycler conditions. The first consisted of 5 µL of 5x MyTaq Red PCR Buffer, 1 µL of 10µM GBT0235ex22 FP2, 1 µL of 10µM GBT0235 WT RP, 0.25 µL of MyTaq DNA Polymerase, 3.5 µL of template, and 14.75 µL dH2O. The second PCR setup replaced the 10µM GBT0235 WT RP with RLC_mRFP R1. Samples were run in a thermal cycler with following PCR conditions: (1) 95°C, 5 min; (2) 95°C, 1 min; (3) 61.3°C, 30 sec; (4) 72°C, 1 min; (5) Go to step 2 39X; (6) 72°C, 5 min; (7) 12°C, hold. Samples were run on a 1% agarose gel and analyzed. We tested our locus-specific *lrpprc* primers (Supplementary Table 1) and found successful amplification of both our *lrpprc*^*+/+*^ and *lrpprc*^*GBT0235/GBT0235*^ bands.

### Genotyping of zebrafish larvae by fin clipping

This larval genotyping protocol was adapted from previously described study (Wilkinson et al., 2017). Larvae were placed in a petri dish containing 3 µL of 20% Tween per 30 mL of embryo media. Larvae were individually placed on an inverted petri dish under a dissecting scope using trans-illumination. A scalpel was used to cut the tail distal to the blood flow. The fin clip was removed from the solution using forceps and placed into a PCR strip tube containing 10 µL of 50 mM NaOH. Scalpel and forceps were sterilized with an ethanol wipe between biopsies. Larvae were transferred into a petri dish with fresh embryo media before being moved to a 24-well plate for recovery while genotyping was performed. Tail biopsies were capped and heated to 98°C for 10 minutes and then cooled to 25°C. 10µL of 100mM Tris pH8 was added to each tube and mixed. These samples were then used for PCR amplification. Following genotyping, embryos were grouped by genotype and placed in petri dishes until needed for experiments.

### Capturing the spatio-temporal expression dynamics in the *in vivo* protein trap lines

Larvae were treated with 0.2 mM phenylthiocarbamide (PTU) at 24 hours post fertilization (hpf) to prevent pigmentation. Fish were anesthetized with 1X tricaine and mounted in 1.5% agarose (Fisher Scientific, USA) prepared with 1X tricaine solution (0.18 mg/L) in an agarose column in the imaging chamber. For Lightsheet microscopy, larval zebrafish were anesthetized with 1X tricaine in embryo water during imaging procedure. RFP expression patterns of 6 dpf larval zebrafish were captured using LP 560 nm filter as excitation and LP 585nm as emission using a Lightsheet Z.1 microscope (Zeiss, Germany) 5X/0.16 NA dry objective. Confocal microscopy was carried out embedding the 6 dpf larvae in 1 % (w/v in E2 medium containing tricaine) low-melting agarose (Fisher Scientific, USA) in a 50-mm glass bottom dish (MatTek, USA). Once the agarose set, it was immersed in E2 medium containing tricaine to prevent drying. Each set of brightfield and RFP images for both heterozygous and homozygous mutants were acquired using Zeiss LSM780 (40XW/1.2 NA) and the images shown are the maximum image projections of the z-stacks obtained from each direction.

### Colocalization analysis

*lrpprc*^*GBT0235/+*^ adult zebrafish were out-crossed to *Tg(MLS: EGFP)* zebrafish (Kim et al., 2008; Sabharwal et al., 2019b) to obtain *Tg(MLS:EGFP) lrpprc* ^*GBT0235/+*^. Embryos were maintained in 0.2 mM phenylthiocarbamide from 24 hpf and anesthetized with tricaine (0.18 mg/L). Larvae sample preparation was done as described above. The 2 dpf larvae were viewed under a Zeiss LSM780 confocal microscope. Laterally oriented Z-stacks of representative images of the caudal fin for GFP and mRFP expressions were captured using the laser with Ex488 and Ex561, respectively under a 63XW/1.2 NA water objective. Each set of images were independently acquired at different fluorescent wavelengths and a composite image was generated using Zeiss Zen microscope software.

### Survival assay

*lrpprc* ^*GBT0235/+*^ adult zebrafish were crossed with *Tg(*−*2.8fabp10:Cre;* −*0.8cryaa:Venus)*^*S955*^ to obtain double transgenic adult zebrafish expressing both the GBT cassette and the fabp10 driven Cre recombinase (liver rescued). At 5 dpf the larval zebrafish were anesthetized using 1X tricaine (0.18 mg/L) and screened for RFP fluorescence and gamma crystalline Venus (V) and sorted. The resultant groups were {RFP-,V- (*lrpprc*^*+/+*^), {RFP-,V+ (*lrpprc*^*+/+*^, liver (fabp10-cre) rescued)}, {RFP+,V+ (*lrpprc*^*GBT0235/+*^ and *lrpprc*^*GBT0235/GBT0235*^, liver (fabp10-cre) rescued)}, {RFP+,V-, BL+(*lrpprc*^*GBT0235/GBT0235*^)}, {RFP+,V-, BL-(*lrpprc*^*GBT0235/+*^). At 5 dpf, the RFP+/V-group was anesthetized and sorted based on the presence of darkened liver (DL) phenotype on a brightfield microscope. The resultant groups were R+/V-/DL+ and R+/V-/DL-. It is assumed at this point that the R+/V- BL+ larvae are the GBT0235 (*lrpprc*) homozygous offspring and the R+/V- BL- larvae are the GBT0235 heterozygous offspring, based on phenotypic characterization of homozygous GBT0235 larval zebrafish. The above-mentioned groups were placed on the Mayo Clinic Zebrafish Facility at 6 dpf. Live counts were recorded daily from 6 dpf to 12 dpf. At the end point (12 dpf) the remaining larvae from each group were counted and euthanized for genotyping using NaOH extraction followed by PCR using primers listed in Supplementary Table 1.

### RNA extraction and sample preparation

*lrpprc* ^*GBT0235/+*^ adult zebrafish were incrossed and embryos collected and placed in a 28°C incubator until larvae were ready for RNA isolation. Larvae were sorted for RFP expression on 6 dpf and separated into dishes. RNA isolation was performed using trizol extraction. Briefly, larvae were placed in 500 µL of trizol and homogenized using a handheld homogenizer on ice and then incubated at room temperature for 5 minutes. Chloroform was added followed by a 15-minute incubation on ice before spinning in tabletop centrifuge at 4°C for 15 minutes. The aqueous layer was removed and placed into a separate 1.7mL centrifuge tube, some of which was used to genotype the larvae. The remainder was put through a series of ethanol and DNase I treatments to isolate and purify the RNA. Each embryo was genotyped using the *lrpprc* ex22_F, GBT0235_R1, and mRFP R1 and RNA samples were grouped according to genotype. RNA quantity and quality were assured to meet the standards of the Genewiz RNA-sequencing platform using the spectrophotometer.

### Mitochondrial DNA relative gene expression analysis

Adult *lrpprc*^*GBT0235/+*^ zebrafish were mated and embryos were collected. Larvae were analyzed for RFP expression and darkened liver (DL) phenotype at 6 dpf. RFP positive, DL positive fish were used as the experimental group to measure mtDNA transcript expression against RFP negative, DL negative embryos as a control. For each clutch, 3 fish from each condition were sorted for RNA extraction. After initial sorting, individual larvae were placed in a 1.7mL tube with 350 µL of RLT buffer: BME at 100:1. Embryos were then homogenized using ∼30 steel beads and tissuelyzed at max frequency for 5 minutes. Homogenized samples were then added to Maxtract tubes, carefully avoiding the steel beads and a phenol/chloroform preparation was used to separate out nucleic acid material. RNA was then isolated and purified using the RNeasy Micro Kit (Qiagen, USA) according to manufacturer’s specifications and spectrophotometry was performed for sample quantification. 250 ng of RNA per fish was used for reverse-transcription into cDNA using the Thermo Fisher SuperScript II kit (Thermo Fisher Scientific, USA). Requisite no reverse transcriptase (RT) controls were run parallel to test for genomic DNA contamination. cDNA was diluted 16-fold with deionized water and amplified using the SensiFAST SYBR Lo-ROX kit (Bioline, USA) for qRT-PCR analysis. Each sample was run in triplicate for RT conditions and duplicate for NRT conditions. Process was repeated for four individual clutches of GBT0235 embryos. Wild type and mutant transcripts levels were normalized to eukaryotic translation elongation factor 1 alpha 1, like 1(*eef1a1l1*).

### Birefringence assay

*lrpprc*^*GBT0235/+*^ adult zebrafish were in-crossed to obtain a population of *lrpprc*^*+/+*^, *lrpprc*^*GBT0235/+*^, and *lrpprc*^*GBT0235/GBT0235*^ offspring. Embryos & larvae were maintained in E2 embryo medium at 28.5°C at a density of 30 per dish and kept on a 14-hour light/10-hour dark cycle. At ∼96 hours post-fertilization, 12 larvae per parental pair were anesthetized with tricaine (0.18 mg/L) and embedded as described in the previous section. The larvae were viewed using a Stemi SV11 stereomicroscope (Zeiss, Germany) equipped with a polarized lens underneath the stage and polarized lens on the objective lens. The polarized lens on the objective was rotated until the background went dark. Birefringence images were then acquired using a DSLR camera (Canon, Powershot G10) in RAW mode using an ISO of 200, aperture of f/4.5, and an exposure of 1/15 s. Larvae were subsequently rescued from agarose using ultra-fine forceps, placed into 8-strip tubes, euthanized by tricaine overdose, and lysed for DNA extraction using 30 µL of 50 mM NaOH (incubated at 95°C for 25 minutes, then neutralized with 3 µL of 1 M Tris-HCl (pH = 8)). This neutralized lysate was used for PCR genotyping.

RAW images were converted to 8-bit grayscale TIFFs in Adobe Photoshop. Grayscale TIFF birefringence images were imported into NIH ImageJ/FIJI for further analysis. We outlined an area in each individual fish that corresponded to the birefringence from its body using the wand tool (legacy mode, threshold = 16). The birefringence signal from the otolith was specifically excluded from these measurements. All outlined regions were saved as regions of interest (ROIs). Within these ROIs, measurements for area, mean gray value, min gray value, max gray value, and integrated gray value density were obtained. During data collection and analysis, experimenters were blind to the genotype and RFP expression pattern of the larvae. After unmasking the data, area, mean gray value, and integrated gray value density were utilized to make plots and assessed for statistical differences between *lrpprc*^*+/+*^ and *lrpprc*^*GBT0235/GBT0235*^ animals. Each individual data point represents a single animal. Each parental pair represents a biological replicate. This dataset contains five biological replicates from two separate days.

### RNA sequencing and analyses

RNA library preparations and sequencing reactions were performed at GENEWIZ, LLC. (South Plainfield, NJ, USA). Quantification of RNA samples was carried out using Qubit 2.0 Fluorometer (Thermo Fisher Scientific, USA) and further, the RNA integrity was checked using Agilent Tapestation 4200 (Agilent Technologies, USA). RNA sequencing libraries were prepared using the NEB Next Ultra RNA Library Prep Kit for Illumina using manufacturer’s instructions (NEB, USA). Briefly, mRNAs were initially enriched with oligo(dT) beads. Enriched mRNAs were fragmented for 15 minutes at 94°C. Subsequently, cDNA fragments were synthesised, end repaired and adenylated at 3’ends, and universal adapters were ligated to them, followed by index addition and library enrichment by PCR with limited cycles. The sequencing library was validated on the Agilent TapeStation (Agilent Technologies, USA), and quantified by using Qubit 2.0 Fluorometer (Thermo Fisher Scientific, USA) as well as by quantitative PCR (KAPA Biosystems, USA). The sequencing libraries were clustered on a single lane of a flow cell. After clustering, the flowcell was loaded on the Illumina HiSeq4000 instrument according to manufacturer’s instructions. The samples were sequenced using a 2×150bp Paired End (PE) configuration. Image analysis and base calling were conducted by the HiSeq Control Software (HCS). Raw sequence data (.bcl files) generated from Illumina HiSeq was converted into fastq files and de-multiplexed using Illumina’s bcl2fastq 2.17 software. One mismatch was allowed for index sequence identification.

Paired-end reads of 150 bp length generated for the homozygous mutant and the wild type was trimmed with a base quality cut off of Phred score Q30 using Trimmomatic (Bolger et al., 2014). Filtered reads were pseudoaligned to the zebrafish reference genome Zv10 version. Kallisto (Bray et al., 2016) was used for transcript assembly and calculation of relative expression values across transcripts. Tximport (Soneson et al., 2015) was used to summarize transcript level counts at gene level. Differential expression was computed using DESeq2 as counts per million (CPM) (Love et al., 2014). Genes with a fold change of greater than log_2_0.5 and less than log_2_0.5 were considered to be upregulated and downregulated respectively.

Human orthologs of differentially expressed zebrafish genes were identified from ZFIN (Zebrafish Information Network) and Ensembl. Differentially expressed human orthologs of zebrafish genes were run against the human Mitocarta database to identify upregulated and downregulated mitochondrial genes involved in biological pathways resident in mitochondria (Calvo et al., 2016). Differentially expressed genes were entered in the PANTHER (Mi et al., 2019) database to identify the protein class and biological process of these genes. For the differentially expressed genes, we predicted the enriched or depleted pathways by using a web based integrated analysis platform, Genetrail2 (Stockel et al., 2016). Differentially expressed genes along with their respective calculated scores were fed as the input file. Scores were calculated as per the following formula: Score= {-log_10_(p-value) * log_2_fold change}. Kolmogorov-Smirnov test was used as the statistical test for the gene set enrichment analysis to find enriched/depleted biological pathways from the KEGG pathways, Reactome pathways and Wiki pathways.

### Oil Red O staining

For whole mount staining at the larval stage, 8 dpf larvae were fixed in 10% NBF overnight. The Oil Red O staining procedures were followed as described previously (Kim et al., 2013). Briefly, *lrpprc*^*+/+*^, *lrpprc*^*GBT0235/GBT0235*^ and *Tg(fabp10:Cre)lrprrc*^*GBT0235/GBT0235*^ larvae were rinsed three times (5 minutes each) with 1× PBS/0.5% Tween-20 (PBS-T). After removing PBS-T, larvae were stained with a mixture of 300 µL of 0.5% ORO in 100% isopropyl alcohol and 200 µL of distilled water for 15 minutes. Larvae were then rinsed with 1× PBS-Tween for three times, twice in 60% isopropyl alcohol for 5 minutes each, briefly rinsed in PBS-Tween, fixed in 10% NBF for 10 minutes, and then mounted in glycerol for imaging. All outlined regions were saved as regions of interest (ROIs). Within these ROIs, measurements for area were obtained and plots and statistical analyses were performed. Oil Red O images were captured with a color camera in JPEG format. For analysis, all images were de-identified using the sample function in R (https://www.r-project.org, https://rstudio.com). De-identified JPEG images were then imported into NIH FIJI/ImageJ and converted from RGB color images to 8-bit grayscale images. For each batch of images (corresponding to one day of experiments), an image with dark Oil Red O staining and an image with light Oil Red O staining were respectively used to determine the lower and upper gray values (the threshold) for positive staining. Briefly, the lower edge of the threshold was set as the gray value that most pixels representing melanocytes were darker than, but almost all dark Oil Red O staining was brighter than. Similarly, the upper edge of the threshold was determined to be the gray value where most pixels representing the background outside of the fish were brighter than, but some light Oil Red O staining was darker than. This left a range of 40-45 gray values that encompassed the most Oil Red O staining. To quantify only the Oil Red O staining in the liver, we drew a single region of interest that bordered the liver of the most darkly stained larvae. This region of interest (ROI) specifically excluded the area around the swim bladder due to accumulation of Oil Red O solution surrounding the swim bladder in a majority of images. Within this ROI, the area of pixel values within the threshold in each larva was determined using the “limit to threshold” option in the “set measurements” menu in FIJI. After areas of Oil Red O staining were extracted from the images, they were re-identified into three groups: *lrpprc*^*+/+*^, *lrpprc*^*GBT0235/GBT0235*^, and *Tg(fabp10:Cre)/lrpprc*^*GBT0235/GBT0235*^.

### Electron microscopy of hepatocyte mitochondria and image analysis

Wild-type, *lrpprc*^*GBT0235/GBT0235*^, *and Tg(fabp10:Cre)/lrpprc*^*GBT0235/GBT0235*^ zebrafish larvae were fixed and imaged using transmission electron microscopy as described in the previous study (Wilson et al., 2019). Briefly, 4 dpf embryos were fixed for 1 to 3 hours in a mixture of 3% glutaraldehyde, 1% formaldehyde, and 0.1 M cacodylate. Embryos were embedded in 2% low melt agarose followed by processing as described earlier (Zeituni et al., 2016). Following embedding steps, post-fixation for 1 hour with 1% osmium tetroxide and 1.25% potassium ferricyanide in cacodylate solution occurred. Larvae were rinsed twice with water and incubated in 0.05 maleate pH 6.5 before staining overnight at 4°C in 0.5% uranyl acetate. Following overnight incubation, samples were washed with water and dehydrated using ethanol dilution. Samples were washed with propylene oxide and then incubated with propylene oxide/resin followed by an overnight evaporation period. Finally, larvae were embedded in 100% resin at 55°C overnight and then 70°C for three days. Sectioning was performed on Reichert Ultracut-S (Leica Microsystems, Germany), mounted on mesh grids, and stained with lead citrate. Images were taken using a Phillips Technai-12 electron microscope and 794 Gatan multiscan CCD camera. Images were scored for hepatocytes using classical hallmark of lipid droplets and subsequently were de-identified and scored by genotype for mitochondrial morphology, in particular the structure and integrity of mitochondrial cristae.

### Feeding of fluorescently tagged long-chain fatty acids

Larvae were genotyped for number of copies (0, 1, or 2) of the RP2 insertion into the *lrpprc* locus and presence or absence of the liver-specific CRE construct *(−2.8fabp10:Cre; - 0.8cryaa:Venus*). In each of four experiments, an equal number of 6 dpf larvae of each genotype (18–24) was pooled and arrayed separately in Falcon brand 6-well tissue culture plates for delivery of a fluorescently tagged long-chain fatty acid feed. 4,4-difluoro-5,7-dimethyl-4-bora-3a,4a-diaza-*s*-indacene 3-dodecanoic acid (BODIPY™ FL C12; Thermo Fisher Scientific, USA) (4 µg/mL) was emulsified in a solution of 5% chicken egg yolk liposomes (in embryo medium) as previously described (Carten et al., 2011). Since larvae can vary greatly in amount they ingest, non-metabolizable TopFluor Cholesterol (conc, 23-(dipyrrometheneboron difluoride)-24-norcholesterol; Avanti Lipids, USA) was added to the feed solution to serve as a readout of amount ingested, allowing for normalization of the HPLC output. TopFluor Cholesterol is harder to get into solution compared to BODIPY FL C12: embryo medium and 1% fatty acid-free BSA was prewarmed to 30°C. After 4 hours of feeding in a light-shielded shaking incubator (30°C, 30 rpm), larvae were rinsed in fresh embryo media and screened for ingestion (darkened intestines) under a stereomicroscope. Larvae that did not eat were disincluded from the study. Larvae that ate were moved to tissue culture plates with fresh embryo media for an overnight chase period of 12.5 h in a 28.5°C light-cycling (14 h on, 10 h off) incubator. Pools of 8–13 larvae from each group were snap frozen on dry ice in microcentrifuge tubes and stored dry at −80°C for lipid extraction.

### Sample preparation and HPLC analyses of long-chain fatty acids

Frozen samples of pooled larvae were subject to lipid extraction using a modified Bligh-Dyer procedure. Extractions were dried and resuspended in 50 µL HPLC-grade isopropanol (HPLC injection solvent). Sample components were separated and detected by an HPLC system as described previously (Quinlivan et al., 2017). Chromatographic peak baselines were manually delimited and peak areas were automatically measured (Chromeleon 7.2; Thermo Fisher Scientific). Peak areas per larval equivalent were calculated and normalized to the TopFluor Cholesterol peak (non-metabolizable; added to the fluorescent fatty acid feed). To fit a linear model wherein the outcomes may have possible unknown correlations, generalized estimated equations (gee) was performed on the R platform comparing total triglyceride area as well as individual peaks/groups of peaks. Assessment of normality by visual examination of Q-Q (quantile-quantile) plots confirmed that the standard normal curve is a valid approximation for the distribution of the data. P-values were obtained from the standard normal Z-table.

### Statistical analyses

Plots and statistical analyses including Man-Whitney U test, Sidak’s multiple comparisons test, student’s t-test and construction of heat map were performed in GraphPad Prism 8 (GraphPad software). Pearson’s correlation coefficient and Manders split coefficients were used for pixel intensity spatial correlation analysis using coloc2 plugin of Fiji image processing package. Statistical significance was derived by Costes p test. Kolmogorov-Smirnov test was used as the statistical test for the gene set enrichment analysis on GeneTrail2 webserver. Results of the statistical analyses have been described either in the results section or in the figure legends.

## Results

### Spatiotemporal expression dynamics of GBT0235 tagged by the *in vivo* protein trapping

To generate and identify the *in vivo* protein trap mutant, a GBT mutagenesis screen was conducted as described previously (Ichino et al., 2019). Briefly, genomic DNA and mRNA expression analyses revealed that the *in vivo* protein trap line GBT0235 contained a single RP2.1 integration event within intron 22 (38 exons total) of the *lrpprc* gene on chromosome 13 (Figure 1B). The RP2.1 GBT protein trap cassette overrode the transcriptional splicing machinery of the endogenous *lrpprc* gene, creating an in-frame fusion between the upstream endogenous exons and the start codon-deficient mRFP reporter sequence. This mRFP-fusion resulted in a translation product that was predicted to be truncated from 1461 amino acids to 793 amino acids. RFP expression analysis of GBT0235 during zebrafish embryonic development revealed onset of reporter expression as early as one cell stage that could be traced until late larval stages. For instance, animals heterozygous for the GBT0235 allele (hereafter *lrpprc*^*GBT023/5+*^) exhibit ubiquitous RFP expression throughout the body at 6 dpf with strong expression in liver, gut and muscles (Figure 2A). To estimate transcriptional effects of the GBT vector in GBT0235, qPCR was carried out on the cDNA from homozygous mutants using primers spanning exon 22-23 of the genomic locus—the site of insertion. Endogenous *lrpprc* transcript levels were negligible in homozygous mutants, *lrpprc*^*GBT0235/GBT0235*^, as compared to wild type controls, *lrpprc*^*+/+*^, indicating a nearly complete knockdown of the endogenous gene (Figure 2B).

**Figure 2:**
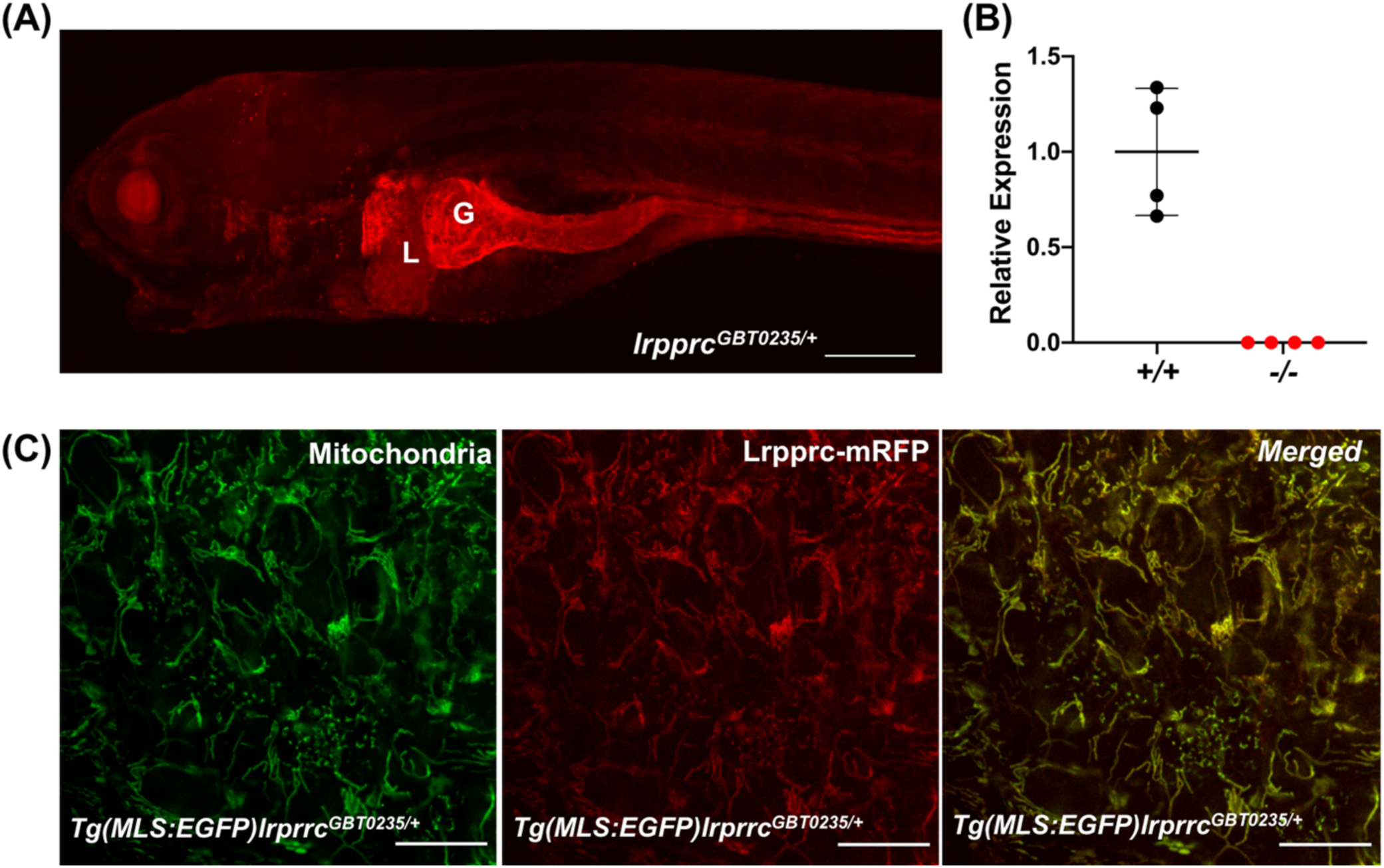
Spatiotemporal expression of Lrpprc-mRFP in GBT0235 mutants. **(A)** Representative images of 6dpf *lrpprc* heterozygous mutants (*lrpprc*^*GBT0235/+*^) with bright RFP expression in the liver and gut (magnification-5X; Scale bar: 200 µm). **(B)** Relative expression of *lrpprc* transcript in wild type (denoted by ‘++”; *lrpprc*^*+/+*^) and homozygous mutant larvae (denoted by “-/-”; *lrpprc*^*GBT0235/GBT0235*^) (p-value <0.001). **(C)** 2 dpf *Tg(MLS:EGFP) lrprrc*^*GBT0235/+*^ was used to observe the sub-cellular localization of the *lrpprc* protein in the caudal fin region. Mitochondria were detected with fluoescence microscopy under GFP filter while the truncated lrprrc:mRFP fusion protein was detected using an RFP filter.

### *lrpprc*^*GBT0235/+*^ mutants exhibit mitochondrial localization of Lrpprc-mRFP protein

In humans *LRPPRC* is a nuclear-encoded gene that encodes for a protein that can be translocated to the mitochondria (Siira et al., 2017; Sterky et al., 2010) where it is known to play an important role in mitochondrial mRNA stability. To investigate the subcellular localization of the mRFP fusion protein (the truncated Lrpprc tagged to mRFP), we crossed *lrpprc*^*GBT0235/+*^ to *Tg(MLS:EGFP)* zebrafish. The *Tg(MLS:EGFP)* line served as a positive control wherein EGFP is fused to the mitochondrial localization signal derived from the zebrafish ortholog of human *COX8A* (Kim et al., 2008). Larvae from the outcross of *lrpprc*^*GBT0235/+*^ and *Tg(MLS:EGFP)* were imaged at high magnification (63X) using confocal microscopy. The caudal fin of 2 dpf (Figure 2C) and myocytes (skeletal muscle region) of 4 dpf (Supplementary Figure 1) zebrafish embryos were selected to observe the mitochondrial network. The caudal fin presents a unique advantage of studying the mitochondrial sub-cellular network *in vivo* as it is as few as two cells thick. Images were taken showing the reticular mitochondrial network marked by the GFP (*Tg(MLS:EGFP)*) and the expression pattern of the Lrpprc-mRFP in skin cells. Resulting images were overlaid to generate a composite image and revealed an overlap between the GFP and Lrpprc-mRFP fusion protein in caudal fin in the zebrafish larvae (Figure 2C). (For caudal fin image: Pearson’s correlation coefficient-0.66; Manders’ split coefficient R1-0.91; Manders’ split coefficient R2-1.0; Costes p-value – 1.0)

### *lrpprc*^*GBT0235/GBT0235*^ mutants develop hallmarks of LSFC

Clinical presentations of patients with LSFC identify a number of hallmarks that we examined in the *lrpprc*^*GBT0235/GBT0235*^ mutants. LSFC patients exhibit metabolic/acidotic crisis followed by death within the first 2 years of life. We therefore hypothesized that our GBT protein trap allele would lead to larval lethality in zebrafish. To assess the survivability, we examined the larvae during the first 12 days of development and obtained a survival curve for the three genotypes (*lrpprc*^*+/+*^, *lrpprc*^*GBT0235/+*^, *and lrpprc*^*GBT0235/GBT0235*^). *lrpprc*^*GBT0235/GBT0235*^ homozygous mutants had a similar survival percentage to that of heterozygous mutants and wild type through 6 dpf. After 6 dpf, mortality was observed in the homozygous mutant group and resulted in 100% lethality at the end of the follow-up on 12 dpf (Figure 3A). In contrast, the survival trend was similar between the heterozygous mutants and the wild type animals. The mortality rate was thus strongly affected by the *lrpprc*^*GBT0235/GBT0235*^ genotype. To assess the impaired function of the *lrpprc* gene, we investigated the transcript levels of mitochondrial-encoded genes. qRT-PCR analysis of these genes revealed a significant drop-off in transcript levels in the *lrpprc*^*GBT0235/GBT0235*^ mutants compared to wild type larvae (Figure 3B). Expression of nuclear genes were found to be similar across the two genotypes. These findings are consistent with data observed in the previous studies that have shown a crucial role for *lrpprc* in mitochondrial mRNA stability.

**Figure 3:**
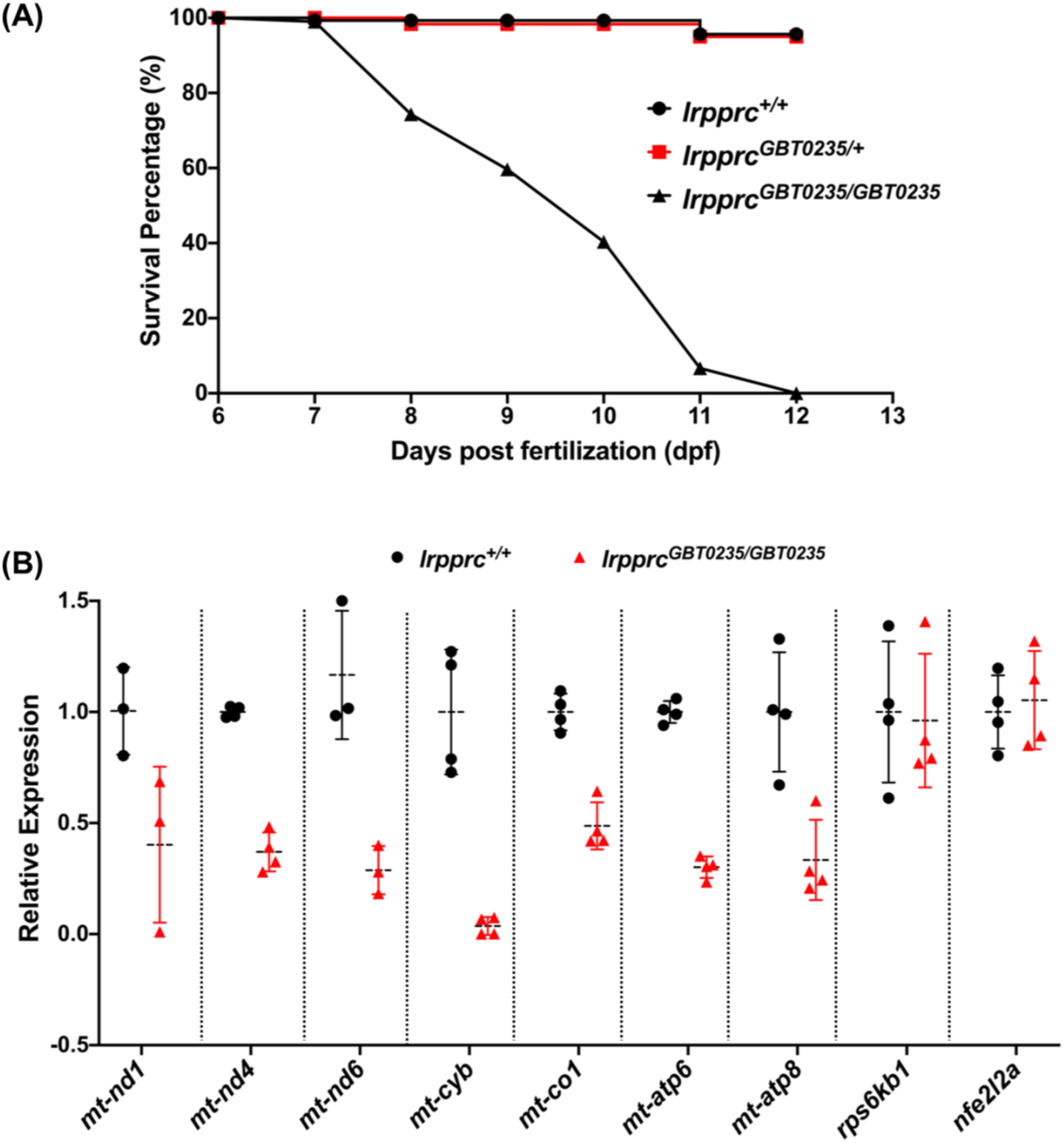
*lrpprc* homozygous zebrafish mutants recapitulate the hallmarks of Leigh syndrome French-Canadian type. **(A)** Survival percentage of three genotypes, wild type *lrpprc*^*+/+*^, heterozygous-*lrpprc*^*GBT0235/+*^ and homozygous mutants-*lrpprc*^*GBT0235/GBT0235*^. Data is represented from independent experiments (p-value <0.05). **(B)** Relative expression of mitochondrial encoded transcripts in the homozygous and wild type siblings assessed by qRT-PCR. Nuclear encoded genes, *rps6kb1 and nfe2l2a* were used as control. Black circle represent wild type and red triangle represent homozygous mutants. Mitochondrial transcripts were normalized to *eef1a1l1* transcript levels. Error bars are represented as SD. (For *mt-nd1*: p-value<0.05, for *mt-nd6, mt-cyb, mt-atp8*: p-value<0.01, for *mt-nd4, mt-co1, mt-atp6*: p-value<0.001, for *rps6kb1* and *nfe2l2a*: p-value –not significant).

Since onset of LSFC has multisystemic involvement, we further investigated for phenotypic defects in tissues such as muscle and liver. Survivors of the early life metabolic crises associated with LSFC often show skeletal muscle phenotypes including hypotonia, muscle weakness, and mobility defects (Sasarman et al., 2015). To investigate whether our *lrpprc*^*GBT0235/GBT0235*^ mutants recapitulate this phenotype, we used birefringence—the refraction of plane polarized light through a complex, ordered structure—as a readout of skeletal muscle development and structure in 4 dpf larvae. *lrpprc*^*GBT0235/GBT0235*^ larvae exemplified (Figure 4A and 4B) a decrease in integrated density and mean gray value when compared with wild type siblings (Figure 4C-4E) and, indicating a deficiency in muscle development specific to *lrpprc*^*GBT0235/GBT0235*^ mutant larvae.

**Figure 4:**
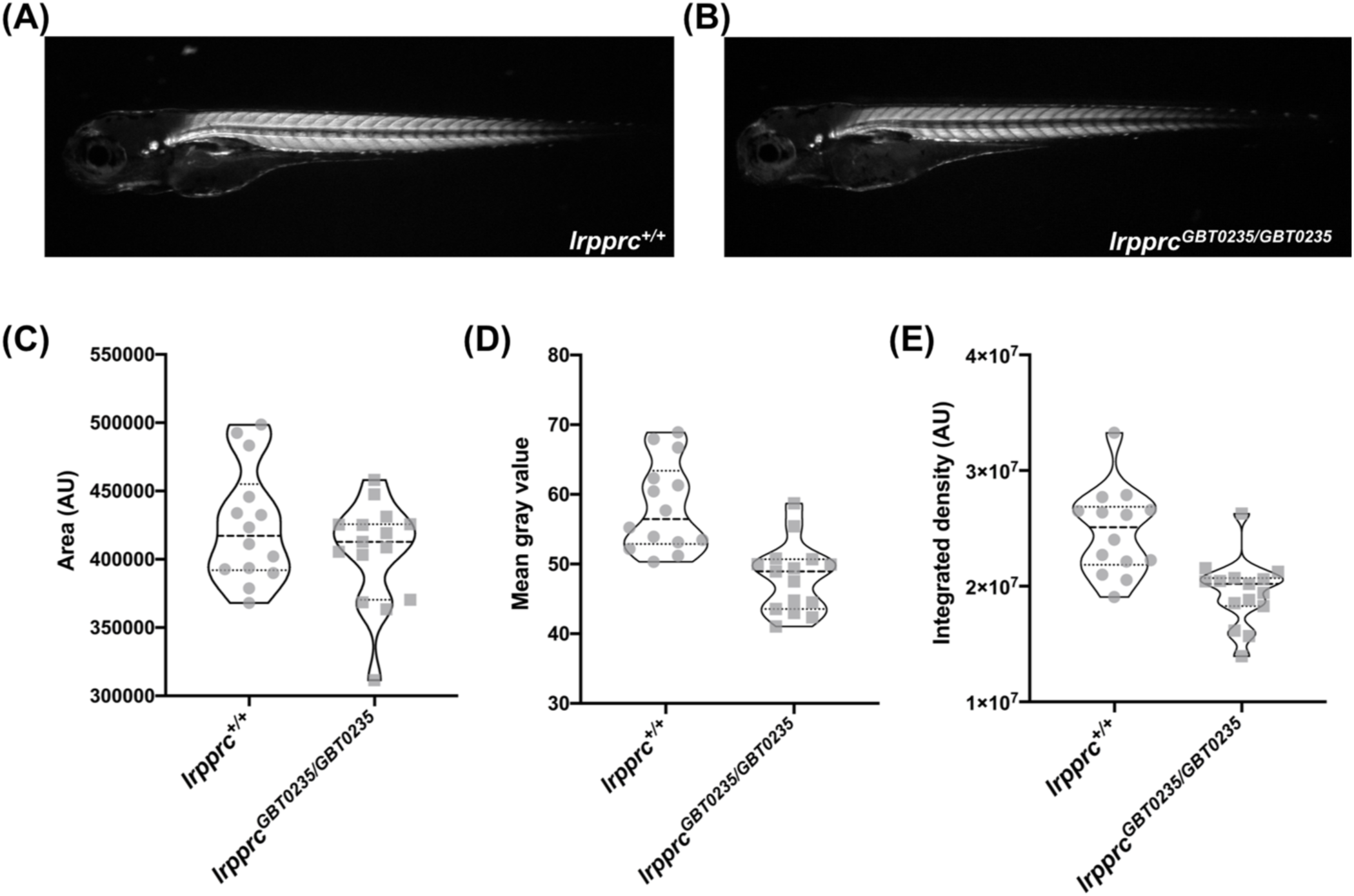
*lrpprc* homozygous mutants display decreased birefringence. **(A-B)** Representative Birefringence images of wildtype **(A)** and *lrpprc*^*GBT0235/GBT0235*^ mutants **(B). (C-E)** The images and graphs in the figure show the area of region of interest (ROI) **(C)**, mean gray value **(D)** and integrated density **(E)** between wild type and *lrpprc*^*GBT0235/GBT0235*^ mutants of the birefringence signal. *lrpprc*^*GBT0235/GBT0235*^ mutants display similar birefringence area but a decrease in mean gray value (p value<0.0001) and integrated density (p value=0.001). Each individual data point represents a single animal. Each parental pair represents biological replicate.

### *lrpprc*^*GBT0235/GBT0235*^ mutants display altered transcriptomic signature

To assess genome wide transcriptional changes in *lrpprc*^*GBT0235/GBT0235*^ mutants, strand specific paired end RNA sequencing (Illumina, USA) with mean read length of 150 bp were generated using HiSeq sequencing platform (Illumina, USA). Approximately 67 million reads were generated for both the *lrpprc*^*+/+*^ and *lrpprc*^*GBT0235/GBT0235*^ RNA preparations. Approximately 88% of the bases had a quality score of >30. The reads were aligned onto the Zv10 reference genome after filtering low quality reads with cut-off Q30 using Trimmomatic software (Bolger et al., 2014). Reads were pseudoaligned using kallisto (Bray et al., 2016) with an average mapping percentage of 79.4%, and approximately 46.18% uniquely mapped reads, and were further assembled across the genome to quantify transcript expression. Transcript level expression was summarized at gene level using Tximport (Soneson et al., 2015). To compare quantitative expression of genes across conditions, we performed differential expression analysis using DESeq2 (Love et al., 2014). The number of transcripts obtained with CPM>1 is as follows: 19205 in WT and 20424 in *lrpprc* homozygous mutant respectively. Empirical cutoff of genes with log_2_Fold Change >= 1 (p-value < 0.1) and log_2_Fold Change <= −1 (p-value < 0.1) were shortlisted as upregulated and downregulated genes respectively (Figure 5A). 825 genes were observed to be significantly upregulated and 776 genes were significantly downregulated in the *lrpprc* homozygous mutants. The human Mitocarta2.0 catalog (Calvo et al., 2016) is a list of 1158 nuclear and mtDNA genes encoding for proteins localized in mitochondria. Overall, 30 and 58 human orthologs of zebrafish upregulated (Supplementary Table 2) and downregulated genes (Supplementary Table 3), respectively overlapped with the database (Figure 5B).

**Figure 5:**
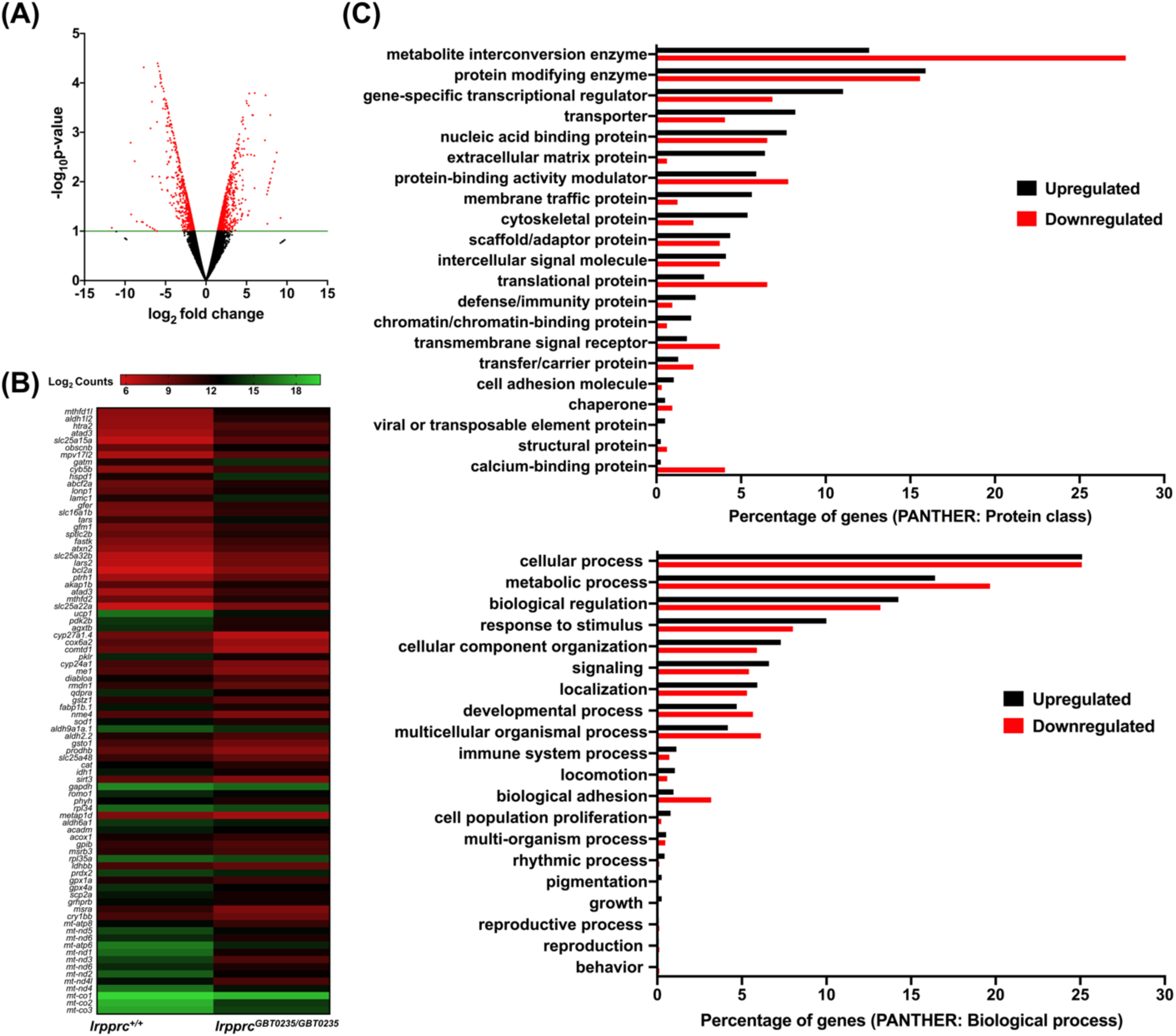
RNAseq of *lrpprc*^*GBT0235/GBT0235*^ homozygous mutants. **(A)** Volcano plot of differentially expressed genes in the homozygous mutants. Logarithm (base 2) of fold change is represented on the x-axis and logarithm of p-value (base 10) is represented on y-axis. Red dot signifies the significantly differentially expressed genes and black dots represent the non-significantly differentially expressed genes between the 6 days post fertilization homozygous *lrpprc*^*GBT0235/GBT0235*^ and wild-type *lrpprc*^*+/+*^ larvae. **(B)** Heat map visualization of expression of zebrafish orthologs for human Mitocarta genes. Gradient color scale represents the log_2_CPM value obtained for each of the zebrafish mitochondrial orthologs in the two data sets. **(C)** PANTHER classification for all the significantly differentially expressed genes in the homozygous mutant according to protein class and biological process. Each histogram represents the percentage of genes falling in each of the category.

Functional classification analysis of the differentially expressed genes using PANTHER (Mi et al., 2019) was performed on two bases: protein class based on the biochemical property and the other was biological process (Figure 5C and Supplementary File 1) in which a gene is involved in the cellular niche. 11% and 12% of the upregulated genes were gene-specific transcriptional regulator and metabolic interconversion enzymes, respectively. Other protein classes that were majorly represented were nucleic acid binding proteins, transporter and protein modifying enzymes. 28% and 16% of the downregulated genes fell in the category of protein class belonging to metabolic interconversion enzyme and protein modifying enzymes. Other protein classes that were majorly represented were translational protein, protein binding activity modulator, nucleic acid binding proteins and gene-specific transcriptional regulator. Most of the differential expressed genes were involved in biological processes such as metabolic process and cellular process and biological regulation. Gene set enrichment analysis predicted mitochondrial respiration and lipid metabolism pathways to be depleted in the *lrpprc*^*GBT0235/GBT0235*^ mutants (Supplementary File 2). These pathways followed a similar trend as observed by the expression of mitochondrial encoded transcriptome and *lrpprc* transcript. Certain signaling pathways such as MAPK signaling, JAK-Stat signaling were also found to be enriched in these animals.

### *lrpprc*^*GBT0235/GBT0235*^ mutants show altered lipid metabolism

Liver is a tissue that has high energetic demand and thus mitochondria are abundant in hepatocytes. The spectrum of the expression of the Lrpprc-mRFP protein was well captured in the liver of the heterozygous (Figure 6A) and homozygous zebrafish mutants (Figure 6B), revealing visible hepatomegaly visible at the gross morphology level in larval animals. Interestingly, *lrpprc*^*GBT0235/GBT0235*^ mutants further display a darkened liver phenotype under brightfield microscopy, reinforcing the involvement of the liver as distinguishing feature in LSFC as compared to traditional Leigh Syndrome (Figure 6C). This hepatomegaly and higher contrast liver phenotype were used as the basis to investigate what role might be played by the liver in this disease. First, we used Oil Red O staining to investigate whether there was a change in the distribution of lipids in *lrpprc*^*GBT0235/GBT0235*^ mutants. We found increased lipid accumulation in the livers of homozygous mutants (Figure 6E and 6G) compared to wild type (Figure 6D and 6G), suggesting the observed darkened liver phenotype could be caused by a change in the density of the stored lipids.

**Figure 6:**
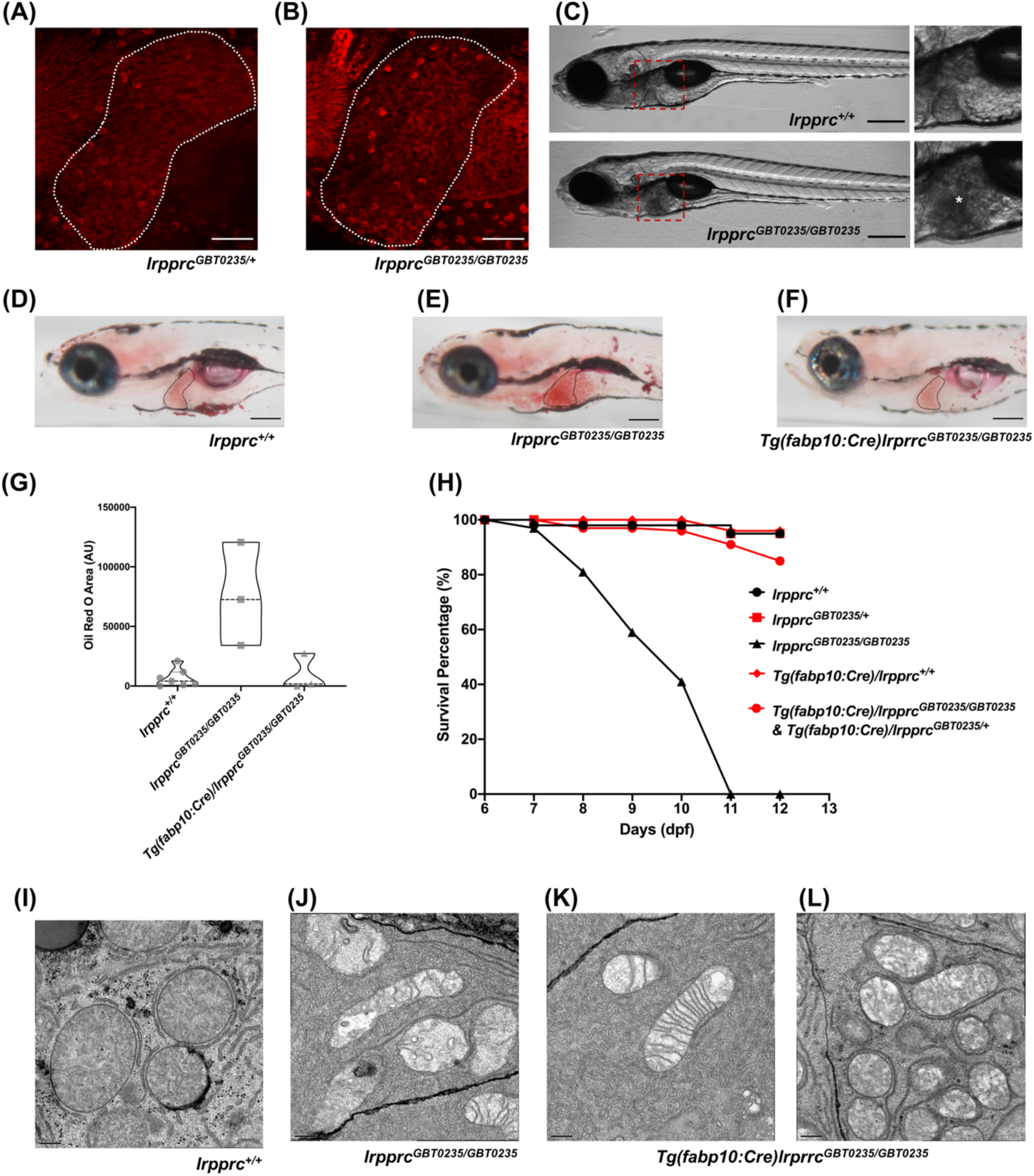
Liver plays an important role in the pathology of LSFC and genetic liver specific rescue rescues the lipid defect and mortality in *lrpprc* homozygous mutant larvae. **(A-B)** Representative images of the livers of 6 dpf old heterozygous (A) *lrpprc*^*GBT0235/+*^and homozygous (B) *lrpprc*^*GBT0235/GBT0235*^ mutants at 40X (Scale bar: 50 µm). **(C)** Brightfield image of 6 dpf wild type, *lrpprc*^*+/+*^ and homozygous *lrpprc*^*GBT0235/GBT0235*^ mutants. Homozygous mutants display dark liver phenotype as compared to the wild type controls. Region showing dark liver has been marked by asterisk. **(D-F)** Oil red O staining for assessment of lipid accumulation in the 8 dpf mutants and 8 dpf rescued larvae. Increased lipid accumulation was observed in the homozygous mutants **(E)** compared to wild type larvae **(D)**. In the liver-specific rescued homozygous *lrpprc* mutants *Tg(fabp10:Cre)lrprrc*^*GBT0235/GBT0235*^, no accumulation of lipids was observed **(F). (G)** The graph show increase in the area of region of interest (ROI) for the accumulated lipids, between wild type and *lrpprc*^*GBT0235/GBT0235*^ mutants, indicating a decrease in the lipid content (p value < 0.05) (Scale bar; G-I: 20 px). The levels are restored in homzygous rescued larvae (p value < 0.05). **(H)** Liver-specific rescued mutants display an improved survival rate beyond 11 dpf. **(I-L)** Representative electron micrographs of the mitochondria in hepatocytes for 8dpf lrpprc wild type **(I)**, *lrpprc* homozygous mutants (J) and liver-specific *lrpprc* rescued larvae **(K and L)**. Altered mitochondrial morphology displayed by *lrrprc*^*GBT0235/GBT0235*^ (J) is improved in the rescued mutants, *Tg(fabp10:Cre)lrprrc*^*GBT0235/GBT0235*^ (K and L) (Scale bar; A-C: 0.5 µm).

To ascertain the hepatic mitochondrial stress which leads to larval lethality (Figure 6H), electron microscopy of the hepatocytes was conducted both for heterozygous and homozygous mutants. Altered morphology of the mitochondria along with irregular shape and distribution of cristae was observed in the hepatocytes of the *lrpprc* homozygous mutants (Figure 6I and 6J).

Since lipid metabolism is a basic liver function dependent on mitochondrial activity, and disruption may lead to acute metabolic crises and death, we further examined how lipid profiles are altered in the null mutants using earlier described methods (Quinlivan et al., 2017) Quinlivan (2017). By coupling the feeding of a fluorescent fatty acid analog with high-performance liquid chromatography methods, we asked whether the *lrpprc* null mutants have differences in fatty acid metabolism compared to wildtype. For these experiments, we fed 6 dpf larvae for 4 hours in a 5% chicken egg yolk solution containing a fluorescent long-chain fatty acid analog (BODIPY™ FL C12) and a non-metabolizable fluorescent reagent (TopFluor^®^ Cholesterol) to assess and correct for amount ingested. After a 12.5-hour chase in fresh media, we extracted total lipids and subjected them to HPLC with fluorescent detection. Extracted lipids were injected across a nonpolar column which separates lipid species by solubility. While chromatograph peak areas were lower overall in the homozygotes (Figure 7A), indicating less ingestion, the non-metabolizable fluorescence allows us to correct for differences (Figure 7B). After chromatograph peak areas were normalized to the non-metabolizable reagent, we found that the *lrpprc*^*+/+*^, *lrpprc*^*GBT0235/+*^, and *lrpprc*^*GBT0235/GBT0235*^ larvae channeled the dietary fluorescent FA into cholesterol ester in similar levels, allowing for efficient cholesterol transport. However, we found that the null mutants incorporated twice as much of the dietary fluorescent FA into non-polar lipids compared to their wildtype siblings (*lrpprc*^*GBT0235/GBT0235*^*/lrpprc*^*+/+*^ = 2.040, 95% CI = 1.122–3.709, p= 0.019; (Figure 7C)).

**Figure 7:**
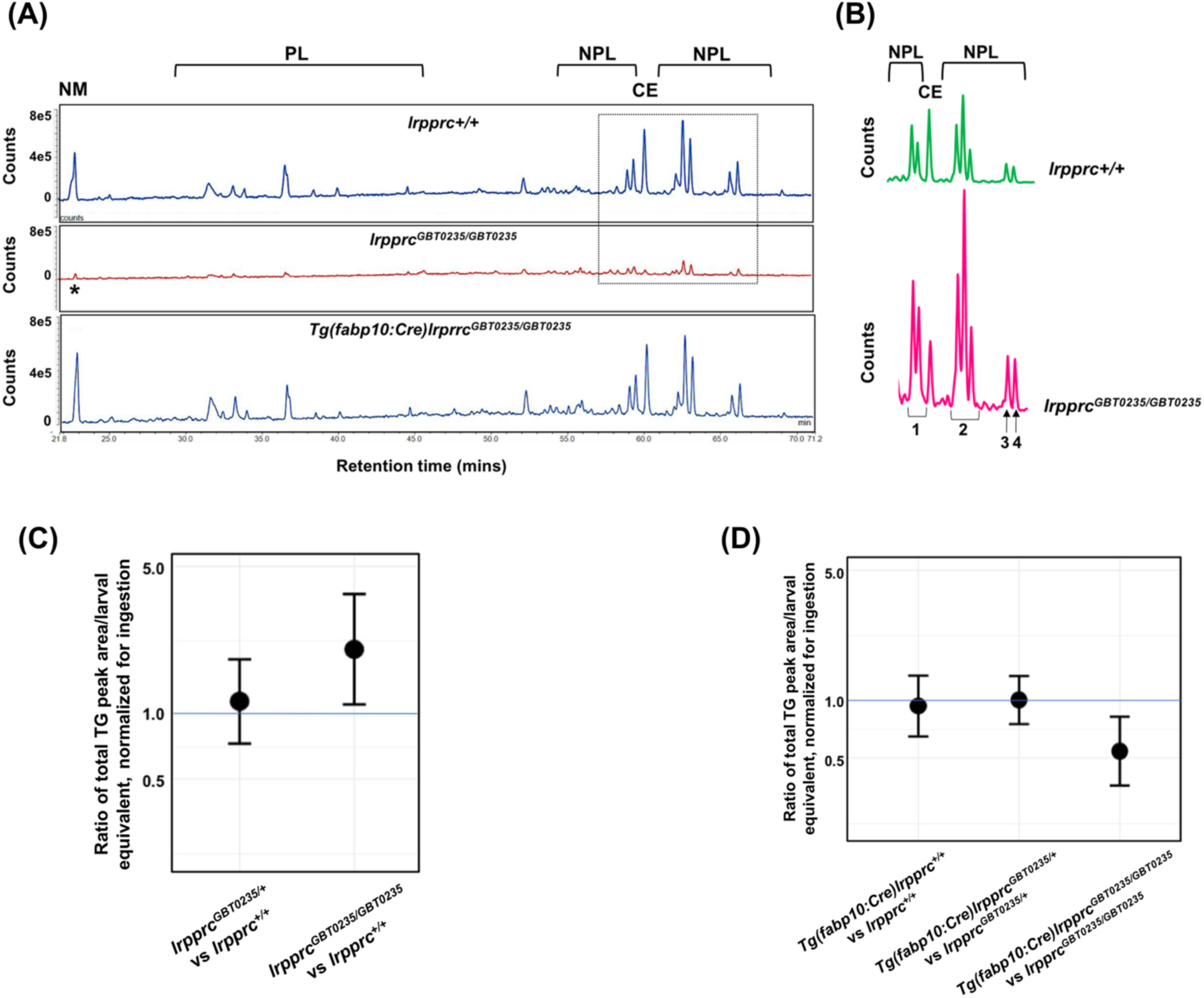
Genetic liver-specific rescue of altered dietary lipid metabolism in *lrpprc* homozygous mutant larvae. **(A-D)** HPLC studies reveal that *lrpprc* homozygous mutants have double the level of nonpolar lipids compared to their wildtype siblings. **(A)** Representative chromatographs are shown: lipid extractions from whole wildtype larvae (top) and whole *lrpprc* homzygous mutant larvae (middle). Peaks represent different lipid species and are quantified by peak area. Measuring the peak of the non-metabolizable fluorescent reagent (22–23 min elution time) confirms that the *lrpprc*^*GBT0235/GBT0235*^ mutants ingest less than their wildtype siblings and provides a normalizer for the other measured peaks. Liver-specific rescue of *lrpprc*^*GBT0235/GBT0235*^ mutants restores nonpolar lipid levels to wildtype levels liver-specific rescued homozygous mutant larvae (bottom). All peaks were normalized to the peak marked by asterisk(*) reflecting non-metabolizable fluorescent reagent (amount ingested). **(B)** Levels of several nonpolar lipid species are higher in *lrpprc* homozygous mutants. Excerpts from representative chromatographs, wildtype larvae (top) and *lrpprc* homozygous mutant larvae (bottom); peaks shown have been normalized to account for amount ingested. Peaks or peak areas (labeled as 1-4) were individually analyzed for contribution to the overall higher NPL level in *lrpprc* homozygous mutants compared to their wildtype siblings. **(C)** 95% CI plot: *lrpprc*^*GBT0235/GBT0235*^ generated 2.04 times more non-polar lipids compared to their wildtype siblings (*lrpprc*^*GBT0235/GBT0235*^/*lrpprc*^*+/+*^ = 2.040, 95% CI = 1.122–3.709, p= 0.019). **(D)** 95% CI plot: *Tg(fabp10:Cre)lrprrc*^*GBT0235/GBT0235*^ restored the levels of non-polar lipids as compared to *lrpprc*^*GBT0235/GBT0235*^ homozygous mutants. PL=phospholipids; NPL=nonpolar lipids (triacylglycerides, diacylglycerides); CE=cholesteryl ester; NM=normalizer for amount eaten.

We next asked whether any particular peak or peak cluster contributed to the higher levels of incorporation into non-polar lipids in the homozygous mutants and found that three of four non-polar lipid peaks or peak clusters are significantly higher (*lrpprc*^*GBT0235/GBT0235*^*/ lrpprc*^*+/+*^ for peaks (1), (2), and (4), respectively = 1.73 [1.02–2.94], 1.99 [1.13–3.52], and 2.91 [1.47–5.79]; p = 0.043, 0.017, and 0.002; Figure 7B). While we are unable to define the exact identities of these peaks at this time, these are likely to be largely made of triacylglycerides and diacylglycerides, the major components of nonpolar lipids along with cholesterol esters. The FA composition, as detected by HPLC-CAD, did not differ between the groups (post-normalization for total lipid levels).

### Liver-specific rescue reverses phenotypes seen in *lrpprc*^*GBT0235/GBT0235*^ mutants

The hepatic dysfunction seen in LSFC patients coupled with the lipid accumulation and darkened liver phenotype in GBT0235 mutants pointed to the liver as an important potential therapeutic avenue. The revertible nature of the GBT construct enables the organismal or tissue-specific reversion to wild type by removal of the cassette via Cre recombinase. Therefore, by crossing *lrpprc* mutants to *Tg(*−*2.8fabp10:Cre;* −*0.8cryaa:Venus)*^*S955*^ zebrafish, we were able to create a liver-rescued *lrpprc* zebrafish, *Tg(fabp10:Cre)/lrpprc*^*GBT0235/GBT0235*^ line that could then be scored for loss of liver-specific mRFP (Supplementary Figure 2) and tested for reversion of the homozygous mutant phenotypes. Following the generation of this line, we replicated the survival experiments to assess the lipid metabolism to determine whether we would see an improvement in liver function. Overall, we noted a decrease in the cholesterol and neutral lipid accumulation in the Oil Red O staining in the liver rescue conditions (Figure 6F and 6G). We also saw a reversion of the lethality in larvae with a mutant allele with 85% of those with the liver rescue surviving until at least 12 dpf (Figure 6H). Liver-specific rescue in the homozygous mutant background reverted the mitochondrial morphology restoring the regular cristae network in hepatocytes (Figure 6K and 6L).

### The altered dietary lipid metabolism in *lrpprc*^*GBT0235/GBT0235*^ mutants is restored to wildtype levels through liver-specific rescue methods

To test the hypothesis that healthy liver function may exert a protective role to overall health in the context of mitochondrial disease, we rescued the liver phenotype of *lrpprc* homozygous mutants by crossing them with a liver-specific Cre recombinase transgenic line and performing an in cross of the double transgenic animals. Since we found altered lipid metabolism in the *lrpprc* null mutants using HPLC methods, we asked whether the liver-specific rescue restores metabolic function to wildtype levels. Not only did the liver-specific rescued null mutants ingest as much as their wildtype and heterozygous siblings, as measured by their non-metabolizable TopFluor Cholesterol peak area, the total peak area of nonpolar lipids were lowered to that of the wildtype siblings (*lrpprc*^*GBT0235/GBT0235*^; *Tg(fabp10:Cre)/lrpprc*^*GBT0235/GBT0235*^ = 0.542; 95% CI = 0.357–0.823, p = 0.004; (Figure 7A)). The genetic background of the liver-specific CRE rescue did not contribute to any change in lipid levels (*Tg(fab10:Cre)lrpprc*^*+/+*^:*lrpprc*^*+/+*^] = 0.944; 95% CI = 0.640–1.391, z = −0.2934, p = 0.769; (Figure 7D)).

## Discussion

In this study, we present a novel zebrafish model for Leigh Syndrome, French Canadian Type (LSFC) identified through the RP2 gene-breaking transposon insertional mutagenesis screen. Phenotypic assessment of this novel allele demonstrates how zebrafish serve as an excellent model to study LSFC. Lrpprc, shares 51% homology and conserved PPR domains with its human ortholog. Bioinformatic analysis starting with the human PPR domains with BLASTP revealed 16 predicted similar PPR domains in the zebrafish *lrpprc* gene indicating the conserved mitochondrial mRNA binding role of this gene in this vertebrate (Supplementary Figure 3). On the basis of location of the RP2 insertion, the RFP protein is predicted to fuse with the N-terminal 793 amino acids of Lrpprc. The spatiotemporal analysis of Lrpprc*-*mRFP shows a co-localization with an established mitochondrial marker in the caudal fin region and myocytes, as the truncated fusion proteins retains the predicted mitochondrial targeting sequence (1-77 amino acids), maintaining its predicted sub-cellular localization. We also observed remarkable ubiquitous expression patterns throughout the organism and hotspots in the liver, skeletal muscle, and eye (Figure 1B). In addition, increased expression of the fusion protein in certain tissues such as the liver, skeletal muscle, and eyes is indicative of the observation that mitochondrial density is tied to energy intensive processes like metabolism, movement, and vision. Different expression data will nevertheless help in expanding our understanding of different roles of this gene in tissue-specific niches.

As the pathophysiology of LSFC is not well understood, we set out to explore the physiological consequences of a *lrpprc* loss-of-function mutation *in vivo* and whether we could mimic the clinical hallmarks seen in LSFC patients. Mutations in *LRPPRC* are associated with infant mortality marked by episodes of fatal acidotic crisis contributing to infant mortality (Morin et al., 1993). Similarly, homozygous *lrpprc* mutant zebrafish displayed complete larval lethality by 12 dpf. However, factors triggering the crisis and associated death in both patients and animal model was initially unclear. Decreased steady state levels of mitochondrially encoded transcripts is another hallmark that is presented in patients with LRPPRC deficiency. Previous studies have implicated LRPPRC to be a critical post-transcriptional regulator of mitochondrial transcripts. LRPPRC regulates the process by the formation of a complex with SRA Stem-Loop Interacting RNA Binding Protein (SLIRP) that recruits mitochondrial polyA polymerase (MTPAP) to initiate the polyadenylation of these transcripts (Ruzzenente et al., 2012). It also acts as a global RNA chaperone stabilizing the mRNA structure and inhibits the 3’ exonucleolytic mRNA degradation regulated by Polynucleotide Phosphorylase (PNPase) and Suppressor of Var1, 3 (SUV3) (Chujo et al., 2012; Siira et al., 2017). Consistent with the previous *in vitro* studies (Gohil et al., 2010), expression of mitochondrial protein coding transcripts was observed to be downregulated in these homozygous mutants by more than 2-fold and were further investigated using RNASeq.

Decreased expression of these protein-coding transcripts is crucial for eliciting defects in the oxidative phosphorylation and mitochondrial respiratory chain. These results were in concordance with the pathways responsible for mitochondrial biogenesis and homeostasis predicted to be depleted in the *lrpprc* homozygous mutants. In addition to the depleted respiratory electron transport and complex 1 biogenesis, bioinformatic analyses also revealed depleted pyruvate metabolism, mitochondrial translation termination and hepatic mitochondrial resident pathway such as metabolism of xenobiotics by cytochrome P450. Various functions including energy production, genome maintenance, ion/metabolite homeostasis, membrane dynamics and transport of biomolecules can be attributed to approximately 1158 nuclear proteins residing in mitochondria (Calvo et al., 2016). We did note an overlap between the set of differentially expressed genes and these 1158 MitoCarta genes, suggesting an altered mitochondrial transcriptome signature. Apart from classical respiratory chain subunits, we identified some key genes from this overlapped data to be differentially expressed that were responsible in pathways such as aldehyde metabolism, fatty acid metabolism and most important, unfolded protein response. Pyruvate dehydrogenase kinase 2b (*pdk2b*) was downregulated in the null mutants and is known to play an important role in the metabolism of fatty acids by inhibiting the pyruvate dehydrogenase activity which inhibits the formation of acetyl-coenzyme A. The expression of mitochondrial isoform of aldehyde dehydrogenase 2 family member, tandem duplicate 2 (*aldh2.2*) was also downregulated, possibly suggesting an interconnection between aldehyde metabolism and detoxification of lipid peroxidation by-products in mitochondria (Nene et al., 2017). *lrpprc* homozygous mutants exhibited an upregulation of chaperones proteins, heat shock protein 60 (*hsp60*) and heat shock protein (*hsp70*) that form an integral part of mitochondrial unfolded protein response. Knockdown of *lrpprc* in a mammalian cell model has been shown to elicit mitochondrial unfolded protein response triggered by nuclear encoded and mitochondrial encoded subunits of complex IV of mitochondrial electron transport chain (Kohler et al., 2015).

Another high energy-requisition tissue on the phenotypic spectrum of LSFC that we explored was muscle. LSFC patients display muscle defects such as hypotonia and COX deficiency (Olahova et al., 2015) and therefore, we adopted a birefringence measurement (Smith et al., 2013) to assess the muscle phenotype in our null mutants. It takes a complex and highly organized structure (like the sarcomeres in skeletal muscle) to rotate the plane polarized light and enable the visualization of birefringence. The noted decrease in birefringence in *lrpprc* mutants is most likely due to the disorganization of sarcomeres and other structures in skeletal muscle. The findings here did concur with those from the transcriptomic studies where key muscle related genes such as myogenic differentiation 1 (*myod1*) and dysferlin (*dysf*) were observed to be differentially expressed. We found *myod1*, a gene involved in muscle regeneration to be downregulated in our study. *dysf* was upregulated in *lrprrc* homozygous mutant larvae, and *Dysf* overexpression was noted in mice to cause muscle defects, including kyphosis and irregular gait (Glover et al., 2010). In addition, we observed the pathway for striated muscle contraction to also show reduced expression in *lrpprc* mutant larvae. Cysteine rich protein 1 b (*csrp1b*), the gene responsible for smooth muscle differentiation and muscle development (Henderson et al., 1999), was also found to be expressed in low levels in the *lrpprc* homozygous mutants.

The liver-specific dysfunction coupled with the episodes of acute metabolic crises distinguishes LSFC from classic Leigh Syndrome (LS). A previous study suggested that liver dysfunction may factor into the increased lactic acidosis and metabolic disturbances as the liver plays a key role in lactate homeostasis via the Cori cycle (Sun et al., 2017). In addition, it is important to note that other basic liver functions that are dependent on mitochondrial activity such as lipid metabolism, serum detoxification, and gluconeogenesis may be key to the development of metabolic crises since they have secondary effects in other organ systems. Under TEM, mitochondrial morphology was altered in the mutants with reduced or collapsed cristae in the hepatocytes, possibly suggesting an impaired mitochondrial function. It has been observed that mitochondrial disorders increase predisposition to intracellular lipid accumulation (Morino et al., 2006). Homozygous *lrpprc* mutants manifested early onset phenotypes with hepatomegaly and progressive accumulation of dark-colored granules in the liver at 6 dpf. These granules appeared to be lipid droplets, which was validated with both Oil Red O staining and extensive *in vivo* feed/chase (i.e., metabolize) labeling lipid studies.

Dietary lipid metabolism studies in the homozygous mutants revealed accumulation of non-polar lipids such as triacylglycerides and diacylglycerides. Accumulation of these lipid species highlights an impaired lipid metabolism in the mutants similar to what is observed in LSFC patients (Ruiz et al., 2019). Metabolic diseases such as nonalcoholic fatty liver disease are associated with accumulation of triglyceride levels and high low-density lipoprotein levels with an underlying mitochondrial dysfunction (Simoes et al., 2018). Hepatic accumulation of such lipids may induce mitochondria to undergo dynamic changes such as depolarization and loss of ATP content (Dominguez-Perez et al., 2019). Further probing of the transcriptomics study revealed that pathways pertaining to fatty acid degradation were depleted in these null mutants. The gene encoding for acyl-coA dehydrogenase medium chain (*Acadm*), a protein that plays an important role in the fatty acid oxidation by making medium-chain acyl-CoA dehydrogenase enzyme was downregulated approximately by fold change of log_2_ 1.5. Catalysis of desaturation of acyl-CoAs to 2-trans-enoyl-CoAs, first step in the fatty acid beta-oxidation pathway, is regulated by acyl-CoA oxidase 1, palmitoyl (*acox1*), expression of which was reduced in mutant larvae. Fatty acid beta-oxidation is not only an exclusive mitochondrial resident pathway but also occurs in peroxisomes. Intrigued to explore that, if the abolishment of *lrpprc* had an inter-organelle effect, we analysed the expression of genes that are involved in peroxisome associated pathways. Surprisingly, MPV17 mitochondrial inner membrane protein like 2 (*mpv17l2*), one of the regulator genes for expression of antioxidant enzymes, was upregulated by fold change of log_2_ 2. Previous study has described that its human ortholog *MPV17L2* downregulates the expression of glutathione peroxidase and catalase genes (Iida et al., 2006). This mechanism was replicated in our mutants where we observed the expression of these genes to be decreased by a fold change of log_2_ 1.7 and log_2_ 1.9 respectively. Overall, our zebrafish LSFC model exhibited a dysfunctional mitochondrial signature both at the transcriptional and phenotypic level, utility of which was explored to design a genetic model therapy for LSFC associated phenotypes.

Because GBT-RP2 contains two loxP sites flanking the protein trap component, we could exploit this utility to rescue the organ specific function of *lrpprc*^*GBT0235/GBT0235*^. To test the hypothesis that liver function might exert a protective role in mitochondrial disease, we induced liver-specific expression of *lrpprc*^*GBT0235/GBT0235*^ by removal of the GBT cassette via liver-specific Cre recombinase. Interestingly, mRFP expression was exclusively extinguished in the liver and reversion of both survival and biochemical phenotypes was observed, suggesting the liver plays a critical role in the pathology of LSFC. Remarkably, these liver-specific rescued larvae exhibited similar levels of non-polar lipids as compared to the wild type larvae, post high-fat meal underscoring the lipid homeostasis as a key therapeutic paradigm in the progression of this disease. Mitigation of abnormal mitochondrial structure with collapsed cristae was also observed in the hepatocytes of the rescued larvae. This effect may in part be due to the restoration of levels of mitochondrial transcripts and thus restoring mitochondrial mediated liver functions such as beta oxidation, fatty acid elongation, cycling of cytochrome molecules and gluconeogenesis. We anticipate that these results, together with those from the conditional knockout experiments will enhance our understanding of the function of *lrpprc* gene in hepatic lipid homeostasis and perhaps shine a light towards new therapeutic options.

With advances in gene editing alongside developments in both viral and non-viral vector delivery, there is hope for tissue-specific gene therapy as a potential avenue to help mitochondrial disease patients. Our findings raise the question whether advances in gene therapy could be used as a therapeutic for patients of LSFC by rescuing *lrpprc* expression in the liver. Since stability of the mitochondrial mRNA transcript, but not its sequence, is disturbed in LSFC, we postulate that inhibiting the mitochondrial RNA degradation pathway may also be a novel therapeutic avenue. Zebrafish are amenable for large-scale drug screens and are increasingly being used for this purpose due to the ease of administration of pharmaceutical compounds. Remarkably, the *lrprrc* homozygous mutants developed various hallmarks of LSFC including decreased mitochondrial transcript stability, aberrant mitochondrial morphology, hepatic steatosis, increased triglyceride levels and early death. Reversion of these measures back to wild type levels could be used as a readout of therapeutic drug candidates.

## Supporting information

Supplementary File 1 PANTHER

Supplementary File 2 PATHWAYS

## Acknowledgements

The authors thank Ms. Mrunal Dehankar of the Mayo Clinic Bioinformatics core for the primary bioinformatics analyses. The authors also thank the Mayo Clinic Zebrafish Facility staff who contributed to the generation of the zebrafish line and the Mayo Clinic Microscopy Cell Analysis and Core for imaging facility. We would also like to acknowledge Mike Sepanski and Vanessa Quinlivan of the Carnegie Institution for the TEM imaging and assistance with HPLC, respectively. This work was funded by NIH grants GM63904 and DA14546, the Marriott Foundation, the Mayo Foundation, R01 DK093399 (Farber, PI). Additional support for this work was provided by the Carnegie Institution for Science Endowment and the G. Harold and Leila Y. Mathers Charitable Foundation. *Tg(*−*2.8fabp10:Cre;* −*0.8cryaa:Venus)*^*S955*^ zebrafish transgenic line was a kind gift from Dr. W. Chen, Vanderbilt University, Nashville, TN, USA. *Tg(MLS:EGFP)* zebrafish transgenic line was a kind gift from Dr. S. Sivasubbu, CSIR-IGIB, India and Dr. S. Choi, Chonnam Medical School, South Korea.

## Author Contributions

The paper was written by AS, MDW, JLA and SCE. Experiments were executed by AS, MDW, RLC, MRS, JLA, AJT, NI, WL, JY, YDing, YDeng with experimental guidance from XX, KJC, SAF and SCE. Data analysis was completed by AS, MDW, RLC, JLA, MRS, AJT, NI, WL.

## Supplementary Figures and Tables

### Supplementary Figures

**Supplementary Figure 1:**
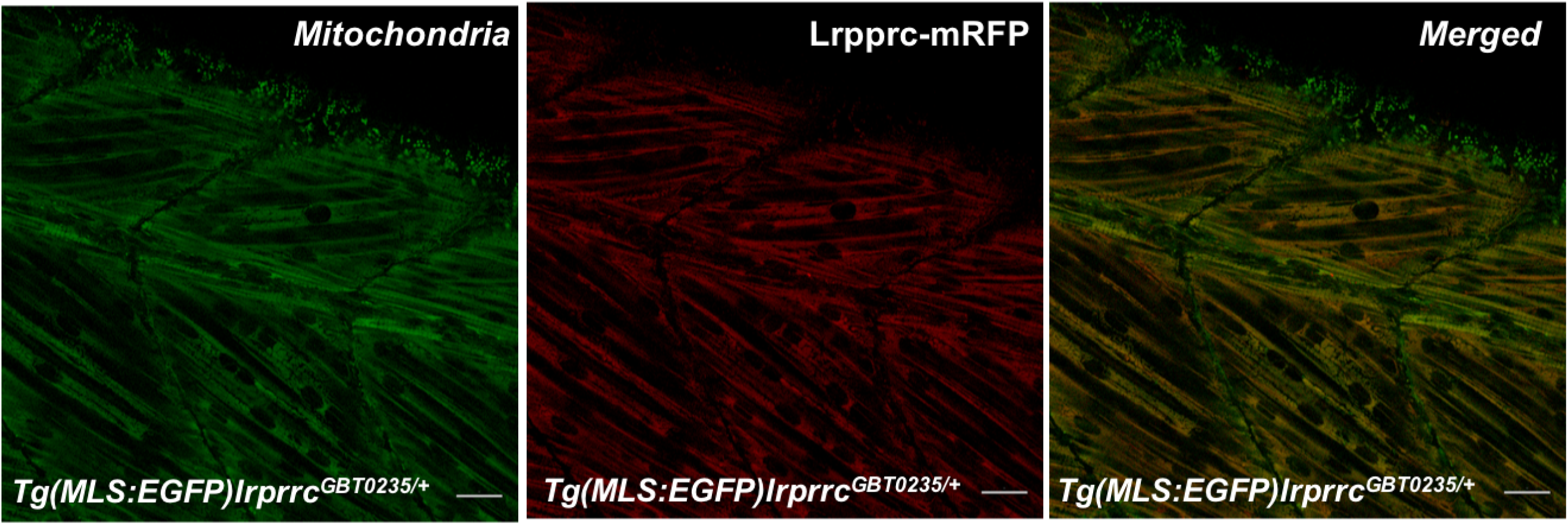
Lrpprc-mRFP localizes to the mitochondria in the zebrafish mutants. 4 dpf *Tg(MLS:EGFP) lrprrc*^*GBT0235/+*^ was used to observe the sub-cellular localization of the Lrpprc-mRFP protein in the myocytes from the skeletal muscle region (Scale bar: 20 µm).

**Supplementary Figure 2:**
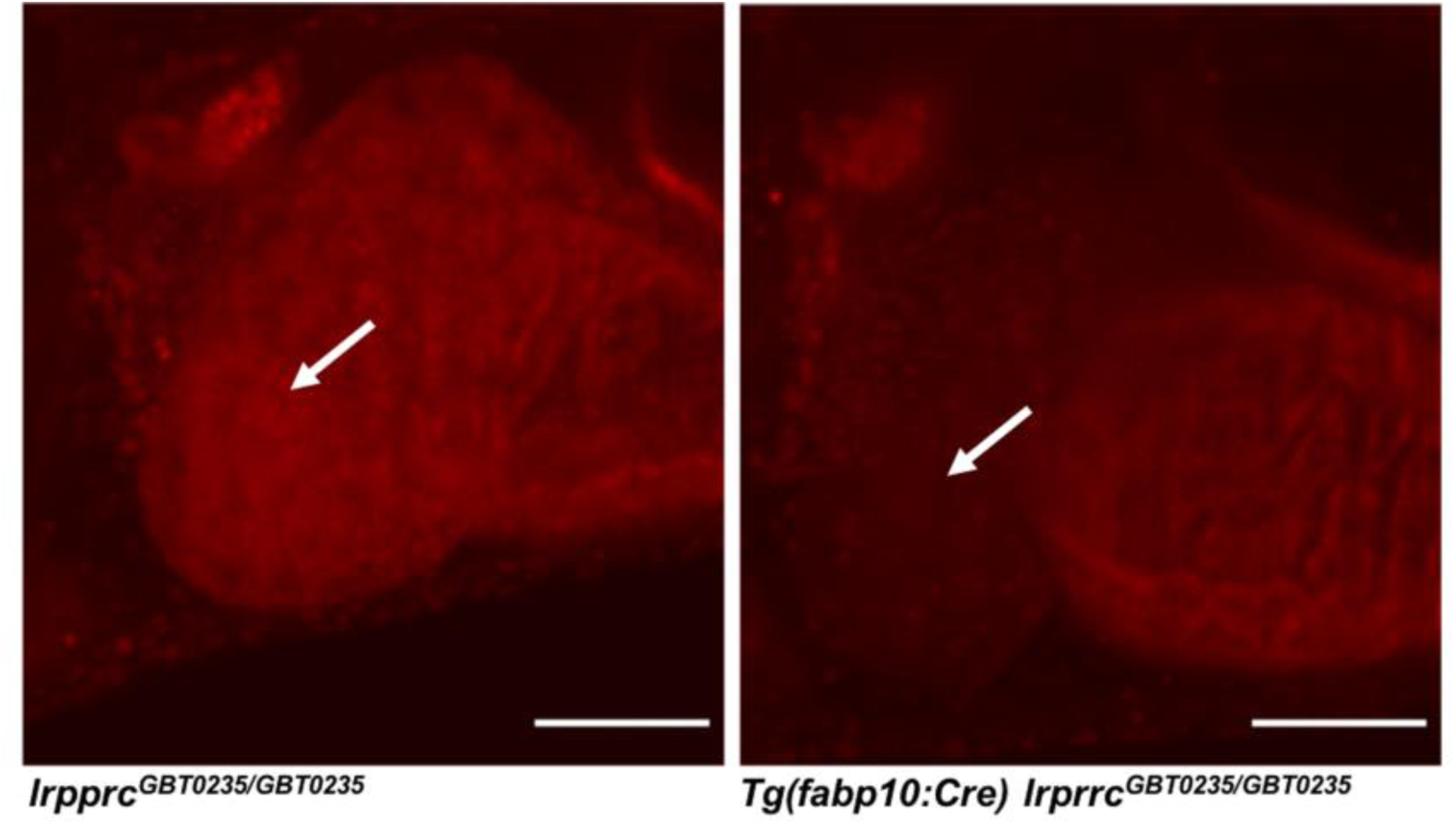
Irreversible liver-specific Cre recombinase mediated rescue. *lrpprc*^*GBT0235/+*^ adult zebrafish was crossed with *Tg(*−*2.8fabp10:Cre;* −*0.8cryaa:Venus)*^*S955*^ to obtain double transgenic adult zebrafish expressing both the GBT cassette and the fabp10 driven Cre recombinase. The liver-specific Cre recombinase rescued the mRFP expression in the liver.

**Supplementary Figure 3:**
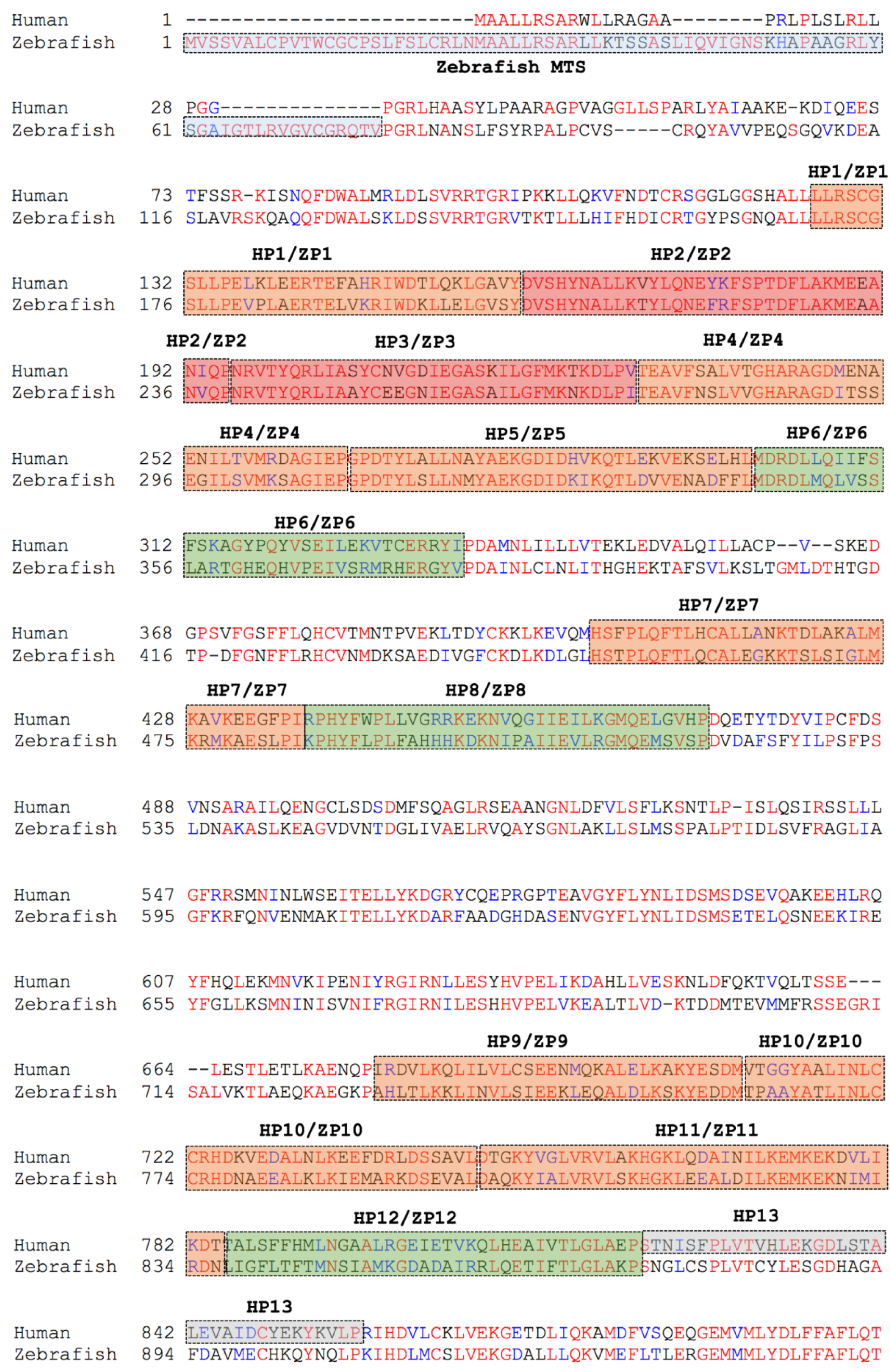
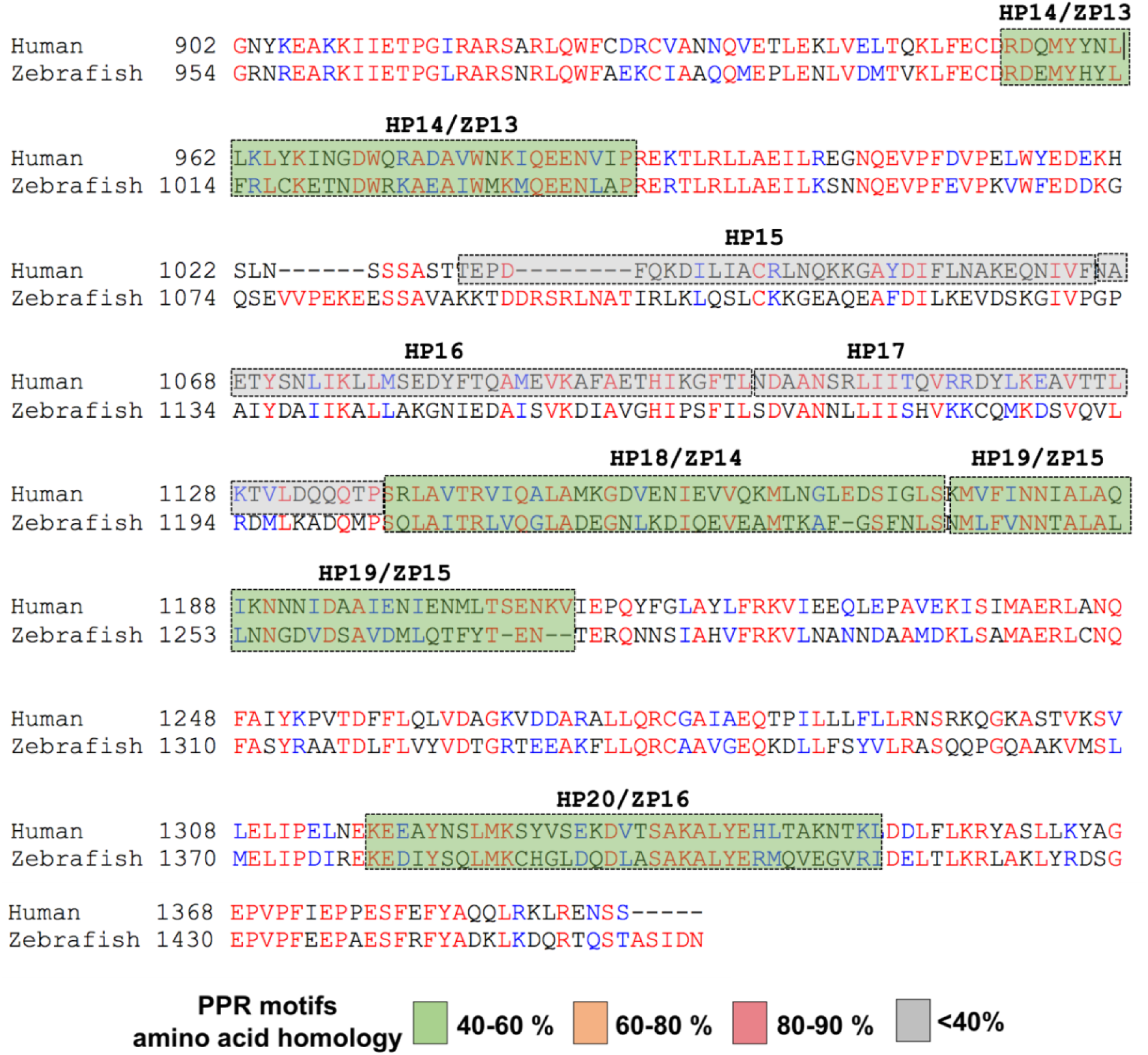
Amino acid alignment of LRRPRC human protein with zebrafish Lrpprc protein. Multiple protein sequence alignment was performed by T-COFFEE multiple sequence alignment webserver. Zebrafish PPR domains were predicted by performing blastp using human PPR domains as the query. The colors of the boxes represent the percentage of homology between the two protein sequences. 16 zebrafish domains were predicted by the homology analysis. Predicted zebrafish mitochondrial targeting sequence is highlighted in blue box (1-77 amino acids). (HP(n): Human PPR domain; ZP(n): Zebrafish PPR domain)

### Supplementary Tables

**Supplementary Table 1:**
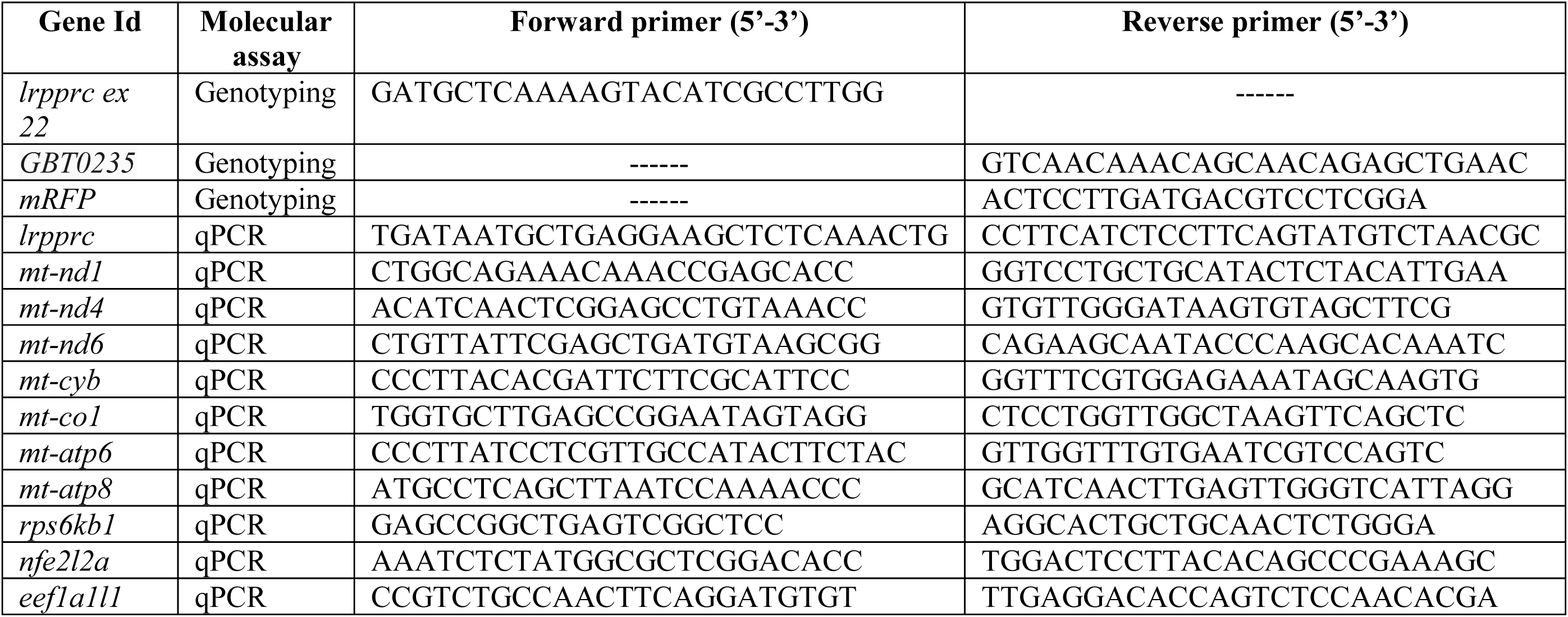
List of oligonucleotides used in the study.

**Supplementary Table 2.**
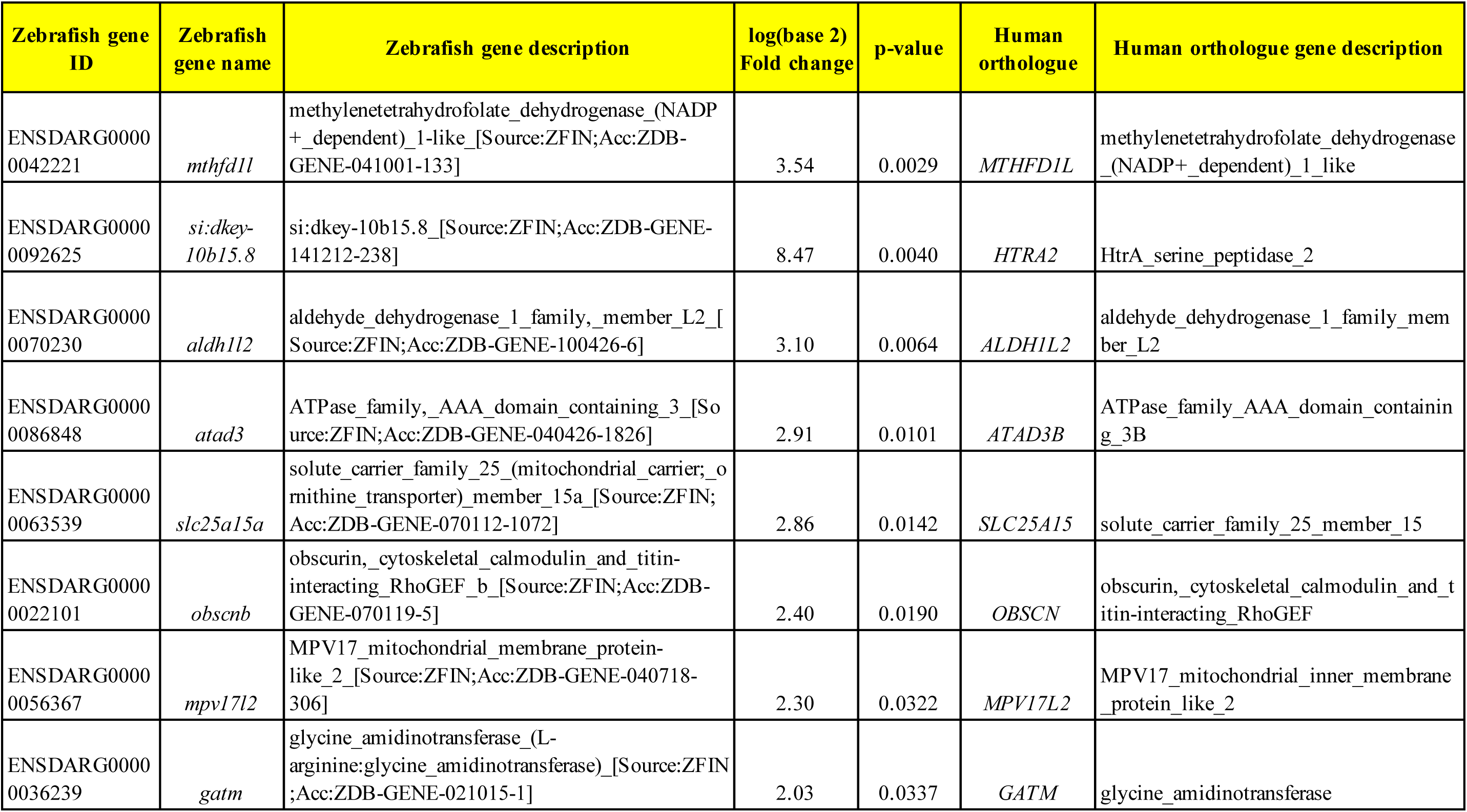
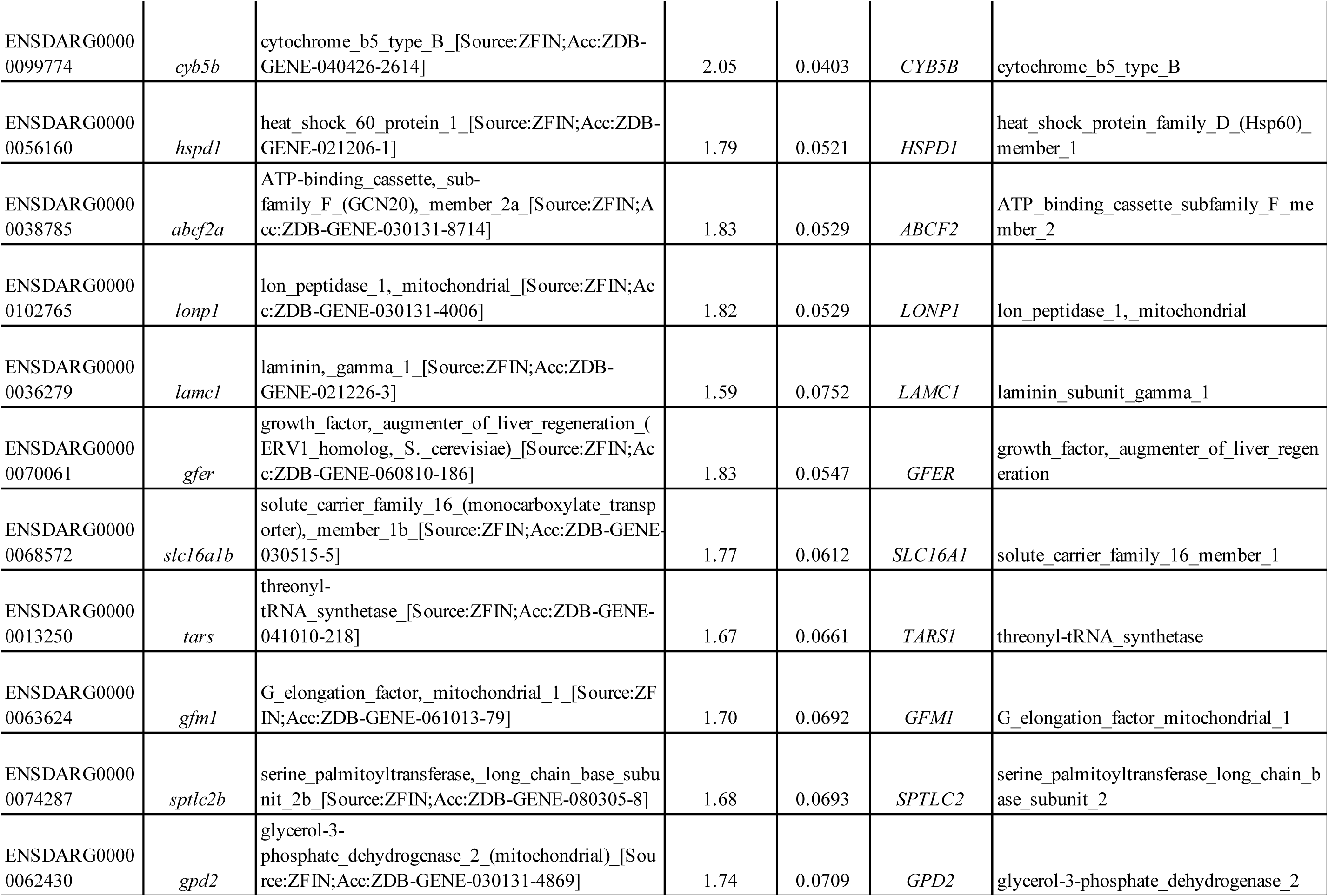
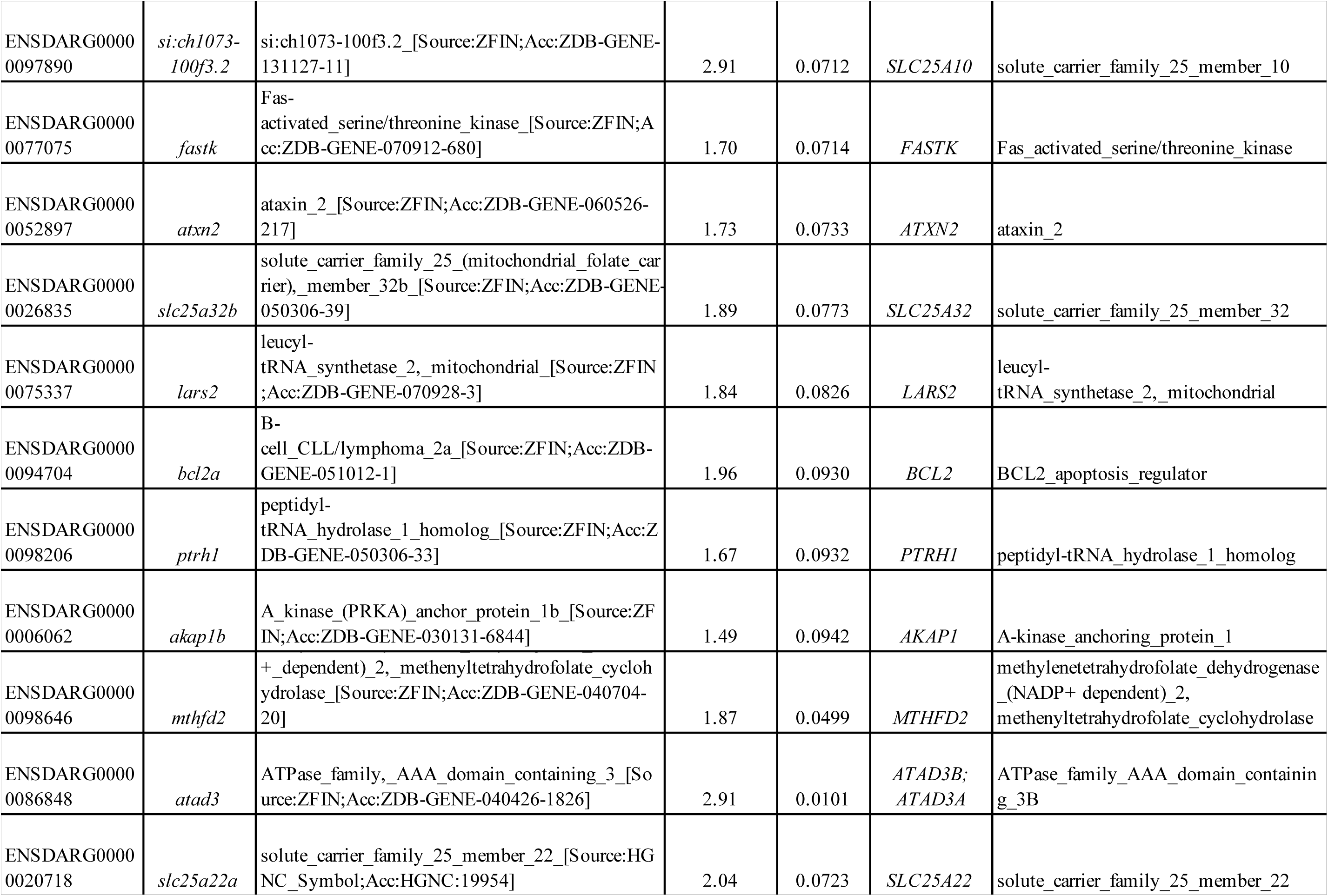
List of human Mitocarta orthologs for zebrafish genes observed to be significantly upregulated in *lrpprc* homozygous zebrafish mutants.

**Supplementary Table 3.** List of human Mitocarta orthologs for zebrafish genes observed to be significantly downregulated in *lrpprc* homozygous zebrafish mutants.

| Zebrafish gene ID | Zebrafish gene name | Zebrafish gene description | log(base 2) Fold change | p-value | Human orthologue | Human orthologue gene description |
| --- | --- | --- | --- | --- | --- | --- |
| ENSDARG00000023151 | *ucpl* | uncoupling_protein_1_[Source:ZFIN;Acc:ZDB-GENE-010503-1] | -3.75 | 0.0017 | *UCPI* | uncoupling protein 1 |
| ENSDARG00000059054 | *pdk2b* | pyruvate_dehydrogenase_kinase_isozyme_2b_[Source:ZFIN;Acc:ZDB-GENE-040426-939] | -3.35 | 0.0035 | *PDK2* | pyruvate dehydrogenase kinase 2 |
| ENSDARG00000052099 | *agxta* | alanine-glyoxylate_aminotransferase_a_[Source:ZFIN;Acc:ZDB-GENE-040718-16] | -3.33 | 0.0038 | *AGXT* | alanine--glyoxylate_and_serine--pyruvate aminotransferase |
| ENSDARG00000055159 | *cyp27a1.4* | cytochrome_P450_family_27_subfamily_A_polypeptide_1_gene_4_[Source:ZFIN;Acc:ZDB-GENE-030131-1060] | -3.12 | 0.0099 | *CYP27A1* | cytochrome_P450_family_27_subfamily A member 1 |
| ENSDARG00000054588 | *cox6a2* | cytochrome_c_oxidase_subunit_VIa_polypeptide_2_[Source:ZFIN;Acc:ZDB-GENE-040912-129] | -2.87 | 0.0111 | *COX6A1* | cytochrome_c_oxidase_subunit_6A1 |
| ENSDARG00000078918 | *comtd1* | catechol-O-methyltransferase_domain_containing_1_[Source:ZFIN;Acc:ZDB-GENE-030131-1072] | -2.72 | 0.0164 | *COMTD1* | catechol-O-methyltransferase_domain_containing 1 |
| ENSDARG00000042010 | *pklr* | pyruvate_kinase_liver_and_RBC_[Source:ZFIN;Acc:ZDB-GENE-010907-1] | -2.44 | 0.0167 | *PKLR* | pyruvate kinase L/R] |
| ENSDARG00000103277 | *cyp24a1* | cytochrome_P450_family_24_subfamily_A_polypeptide_1_[Source:ZFIN;Acc:ZDB-GENE-060825-1] | -2.44 | 0.0204 | *CYP24A1* | cytochrome_P450_family_24_subfamily A member 1 |
| ENSDARG00000053215 | *me1* | malic_enzyme_1,_NADP(+)-dependent,_cytosolic_[Source:ZFIN;Acc:ZDB-GENE-061013-438] | -2.45 | 0.0212 | *ME1* | malic enzyme 1 |
| ENSDARG00000104172 | *diabloa* | diablo,_IAP-binding_mitochondrial_protein_a_[Source:ZFIN;Acc:ZDB-GENE-040426-1303] | -2.30 | 0.0220 | *DIABLO* | diablo_IAP-binding mitochondrial protein] |
| ENSDARG00000074507 | *rmdn1* | regulator_of_microtubule_dynamics_1_[Source:ZFIN;Acc:ZDB-GENE-070928-17] | -2.29 | 0.0241 | *RMDN1* | regulator_of_microtubule_dynamics 1 |
| ENSDARG00000040190 | *qdpra* | quinoid_dihydropteridine_reductase_a_[Source:ZFIN;Acc:ZDB-GENE-070705-197] | -2.21 | 0.0247 | *QDPR* | quinoid_dihydropteridine_reductase |
| ENSDARG00000027984 | *gstz1* | glutathione_S-transferase_zeta_1_[Source:ZFIN;Acc:ZDB-GENE-040718-184] | -2.23 | 0.0258 | *GSTZ1* | glutathione S-transferase zeta 1 |
| ENSDARG00000059227 | *fabp1b.1* | fatty_acid_binding_protein_1b,_tandem_duplicate_1_[Source:ZFIN;Acc:ZDB-GENE-050522-96] | -2.16 | 0.0271 | *FABP1* | fatty acid binding protein 1 |
| ENSDARG00000088390 | *nme4* | NME/NM23_nucleoside_diphosphate_kinase_4_[Source:ZFIN;Acc:ZDB-GENE-040426-1043] | -2.21 | 0.0315 | *NME4* | NME/NM23_nucleoside_diphosphate kinase 4 |
| ENSDARG00000043848 | *sod1* | superoxide_dismutase_1,_soluble_[Source:ZFIN;Acc:ZDB-GENE-990415-258] | -2.03 | 0.0341 | *SOD1* | superoxide dismutase 1 |
| ENSDARG00000069100 | *aldh9a1a.1* | aldehyde_dehydrogenase_9_family,_member_A1a,_tandem_duplicate_1_[Source:ZFIN;Acc:ZDB-GENE-030131-1257] | -2.01 | 0.0346 | *ALDH9A1* | aldehyde_dehydrogenase_9_family_member A1 |
| ENSDARG00000028087 | *aldh2.2* | aldehyde_dehydrogenase_2_family_(mitochondrial),_tandem_duplicate_2_[Source:ZFIN;Acc:ZDB-GENE-030326-5] | -2.01 | 0.0373 | *ALDH2* | aldehyde_dehydrogenase_2_family_member |
| ENSDARG00000022183 | *gstol* | glutathione_S-transferase_omega_1_[Source:ZFIN;Acc:ZDB-GENE-040718-365] | -2.02 | 0.0407 | *GSTO1* | glutathione S-transferase omega 1 |
| ENSDARG00000086512 | *prodhb* | proline_dehydrogenase_(oxidase)_1b_[Source:ZFIN;Acc:ZDB-GENE-101116-1] | -2.10 | 0.0414 | *PRODH* | proline dehydrogenase 1 |
| ENSDARG00000021250 | *slc25a48* | solute_carrier_family_25_member_48_[Source:ZFIN;Acc:ZDB-GENE-040718-60] | -1.97 | 0.0424 | *SLC25A48* | solute_carrier_family_25_member_48 |
| ENSDARG00000104702 | *cat* | catalase_[Source:ZFIN;Acc:ZDB-GENE-000210-20] | -1.90 | 0.0434 | *CAT* | catalase] |
| ENSDARG00000025375 | *idh1* | isocitrate_dehydrogenase_1_(NADP+)_soluble_[Source:ZFIN;Acc:ZDB-GENE-031006-1] | -1.89 | 0.0440 | *IDH1* | isocitrate_dehydrogenase_(NADP(+)) 1, cytosolic |
| ENSDARG00000035819 | *sirt3* | sirtuin_3_[Source:ZFIN;Acc:ZDB-GENE-070112-1762] | -1.93 | 0.0533 | *SIRT3* | sirtuin 3 |
| ENSDARG00000043457 | *gapdh* | glyceraldehyde-3-phosphate_dehydrogenase_[Source:ZFIN;Acc:ZDB-GENE-030115-1] | -1.70 | 0.0597 | *GAPDH* | glyceraldehyde-3-phosphate dehydrogenase |
| ENSDARG00000038076 | *romol* | reactive_oxygen_species_modulator_1_[Source:ZFIN;Acc:ZDB-GENE-040426-1768] | -1.66 | 0.0657 | *ROMO1* | reactive_oxygen_species_modulator 1 |
| ENSDARG00000086740 | *phyh* | phytanoyl-CoA_2-hydroxylase_[Source:ZFIN;Acc:ZDB-GENE-050417-361] | -1.62 | 0.0716 | *PHYH* | phytanoyl-CoA 2-hydroxylase |
| ENSDARG00000029500 | *rpl34* | ribosomal_protein_L34_[Source:ZFIN;Acc:ZDB-GENE-040426-1033] | -1.58 | 0.0747 | *RPL34* | ribosomal protein L34 |
| ENSDARG00000019715 | *metap1d* | methionyl_aminopeptidase_type_1D_(mitochondrial)_[Source:ZFIN;Acc:ZDB-GENE-050522-71] | -1.84 | 0.0799 | *METAP1D* | methionyl_aminopeptidase_type_1D, mitochondrial |
| ENSDARG00000053485 | *aldh6a1* | aldehyde_dehydrogenase_6_family_member_A1_[Source:ZFIN;Acc:ZDB-GENE-030131-9192] | -1.55 | 0.0803 | *ALDH6A1* | aldehyde_dehydrogenase_6_family_member A1 |
| ENSDARG00000038900 | *acadm* | acyl-CoA_dehydrogenase,_C-4_to_C-12_straight_chain_[Source:ZFIN;Acc:ZDB-GENE-040426-1945] | -1.54 | 0.0823 | *ACADM* | acyl-CoA dehydrogenase medium chain |
| ENSDARG00000014727 | *acox1* | acyl-CoA_oxidase_1,_palmitoyl_[Source:ZFIN;Acc:ZDB-GENE-041010-219] | -1.51 | 0.0894 | *ACOX1* | acyl-CoA oxidase 1 |
| ENSDARG00000103826 | *gpib* | glucose-6-phosphate_isomerase_b_[Source:ZFIN;Acc:ZDB-GENE-020513-3] | -1.53 | 0.0899 | *GPI* | glucose-6-phosphate isomerase] |
| ENSDARG00000045658 | *msrb3* | methionine_sulfoxide_reductase_B3_[Source:ZFIN;Acc:ZDB-GENE-040625-74] | -1.48 | 0.0966 | *MSRB3* | methionine sulfoxide reductase B3 |
| ENSDARG00000088030 | *rpl35a* | ribosomal_protein_L35a_[Source:ZFIN;Acc:ZDB-GENE-040718-190] | -1.44 | 0.0970 | *RPL35A* | ribosomal protein L35a[ |
| ENSDARG00000071076 | *ldhbb* | lactate_dehydrogenase_Bb_[Source:ZFIN;Acc:ZDB-GENE-040718-176] | -1.50 | 0.0997 | *LDHB* | lactate dehydrogenase B |
| ENSDARG00000025350 | *prdx2* | peroxiredoxin_2_[Source:ZFIN;Acc:ZDB-GENE-030326-2] | -1.56 | 0.0783 | *PRDX1* | peroxiredoxin 1 |
| ENSDARG00000068478 | *gpx4a* | glutathione_peroxidase_4a_[Source:ZFIN;Acc:ZDB-GENE-030410-2] | -2.02 | 0.0346 | *GPX4* | glutathione peroxidase 4 |
| ENSDARG00000018146 | *gpx1a* | glutathione_peroxidase_1a_[Source:ZFIN;Acc:ZDB-GENE-030410-1] | -1.68 | 0.0668 | *GPX1* | glutathione peroxidase 1 |
| ENSDARG00000012194 | *scp2a* | sterol_carrier_protein_2a_[Source:ZFIN;Acc:ZDB-GENE-040426-1846] | -2.07 | 0.0318 | *SCP2* | sterol carrier protein 2 |
| ENSDARG00000019986 | *grhprb* | glyoxylate_reductase/hydroxypyruvate_reductase_b_[Source:ZFIN;Acc:ZDB-GENE-040426-1847] | -1.52 | 0.0854 | *GRHPR* | glyoxylate and hydroxypyruvate reductase |
| ENSDARG00000054588 | *cox6a2* | cytochrome_c_oxidase_subunit_VIa_polypeptide_2_[Source:ZFIN;Acc:ZDB-GENE-040912-129] | -2.87 | 0.0111 | *COX6A2* | cytochrome_c_oxidase_subunit_6A1_[Source:HGNC_Symbol;Acc:HGNC:2277] |
| ENSDARG00000029587 | *msra* | methionine_sulfoxide_reductase_A_[Source:ZFIN;Acc:ZDB-GENE-041014-344] | -3.19 | 0.0054 | *MSRA* | methionine sulfoxide reductase A |
| ENSDARG00000069074 | *cry1ba* | cryptochrome_circadian_clock_1ba_[Source:ZFIN;Acc:ZDB-GENE-010426-4] | -1.80 | 0.0506 | *CRY1* | cryptochrome circadian regulator 3a |
| ENSDARG00000091131 | *cry1bb* | cryptochrome_circadian_clock_1bb_[Source:ZFIN;Acc:ZDB-GENE-010426-5] | -1.75 | 0.0650 | *CRY1* | cryptochrome circadian regulator 3b |
| ENSDARG00000063895 | *mt-nd1* | NADH_dehydrogenase_1,_mitochondrial_[Source:ZFIN;Acc:ZDB-GENE-011205-7] | -4.63 | 0.0004 | *MT-ND1* | biquinone_oxidoreductase_core_subunit_1_[Source:HGNC_Symbol;Acc:HGNC:7455] |
| ENSDARG00000063899 | *mt-nd2* | NADH_dehydrogenase_2,_mitochondrial_[Source:ZFIN;Acc:ZDB-GENE-011205-8] | -3.91 | 0.0013 | *MT-ND2* | biquinone_oxidoreductase_core_subunit_2_[Source:HGNC_Symbol;Acc:HGNC:7456] |
| ENSDARG00000063914 | *mt-nd3* | NADH_dehydrogenase_3,_mitochondrial_[Source:ZFIN;Acc:ZDB-GENE-011205-9] | -5.21 | 0.0001 | *MT-ND3* | biquinone_oxidoreductase_core_subunit_3_[Source:HGNC_Symbol;Acc:HGNC:7458] |
| ENSDARG00000063917 | *mt-nd4* | NADH_dehydrogenase_4,_mitochondrial_[Source:ZFIN;Acc:ZDB-GENE-011205-10] | -3.68 | 0.0020 | *MT-ND4* | biquinone_oxidoreductase_core_subunit_4_[Source:HGNC_Symbol;Acc:HGNC:7459] |
| ENSDARG00000063916 | *mt-nd4l* | NADH_dehydrogenase_4L,_mitochondrial_[Source:ZFIN;Acc:ZDB-GENE-011205-11] | -3.58 | 0.0024 | *MT ND4L* | NADH_dehydrogenase_4L,_mitochondrial |
| ENSDARG00000063921 | *mt-nd5* | NADH_dehydrogenase_5,_mitochondrial_[Source:ZFIN;Acc:ZDB-GENE-011205-12] | -2.77 | 0.0093 | *MT-ND5* | mitochondrially_encoded_NADH:ubiquinone_oxidoreductase_core_subunit_5 |
| ENSDARG00000063922 | *mt-nd6* | NADH_dehydrogenase_6,_mitochondrial_[Source:ZFIN;Acc:ZDB-GENE-011205-13] | -2.88 | 0.0078 | *MT-ND6* | mitochondrially_encoded_NADH:ubiquinone_oxidoreductase_core_subunit_6 |
| ENSDARG00000063924 | *mt-cyb* | cytochrome_b_mitochondrial_[Source:ZFIN;Acc:ZDB-GENE-011205-17] | -4.88 | 0.0003 | *MT-CYB* | mitochondrially_encoded_cytochrome b |
| ENSDARG00000063905 | *mt-col* | cytochrome_c_oxidase_I_mitochondrial_[Source:ZFIN;Acc:ZDB-GENE-011205-14] | -1.61 | 0.0710 | *MT-COI* | mitochondrially_encoded_cytochrome c oxidase I |
| ENSDARG00000063908 | *mt-co2* | cytochrome_c_oxidase_II_mitochondrial_[Source:ZFIN;Acc:ZDB-GENE-011205-15] | -4.59 | 0.0004 | *MT-CO2* | mitochondrially_encoded_cytochrome c oxidase II |
| ENSDARG00000063912 | *mt-co3* | cytochrome_c_oxidase_III_mitochondrial_[Source:ZFIN;Acc:ZDB-GENE-011205-16] | -3.91 | 0.0013 | *MT-CO3* | mitochondrially_encoded_cytochrome c oxidase III |
| ENSDARG00000063911 | *mt-atp6* | ATP_synthase_6_mitochondrial_[Source:ZFIN;Acc:ZDB-GENE-011205-18] | -3.51 | 0.0026 | *MT-ATP6* | mitochondrially_encoded_ATP_synthase membrane subunit 6 |
| ENSDARG00000063910 | *mt-atp8* | ATP_synthase_8_mitochondrial_[Source:ZFIN;Acc:ZDB-GENE-011205-19] | -2.69 | 0.0109 | *MT-ATP8* | ATP synthase 8, mitochondrial |

## References

Bolger AM, Lohse M, Usadel B. 2014. Trimmomatic: a flexible trimmer for Illumina sequence data. Bioinformatics 30:2114–2120. doi:10.1093/bioinformatics/btu170

Bray NL, Pimentel H, Melsted P, Pachter L. 2016. Near-optimal probabilistic RNA-seq quantification. Nat Biotechnol 34:525–527. doi:10.1038/nbt.3519

Broughton RE, Milam JE, Roe BA. 2001. The complete sequence of the zebrafish (Danio rerio) mitochondrial genome and evolutionary patterns in vertebrate mitochondrial DNA. Genome Res. doi:10.1101/gr.156801

Byrnes J, Ganetzky R, Lightfoot R, Tzeng M, Nakamaru-Ogiso E, Seiler C, Falk MJ. 2018. Pharmacologic modeling of primary mitochondrial respiratory chain dysfunction in zebrafish. Neurochem Int. doi:10.1016/j.neuint.2017.07.008

Calvo SE, Clauser KR, Mootha VK. 2016. MitoCarta2.0: an updated inventory of mammalian mitochondrial proteins. Nucleic Acids Res 44:D1251–7. doi:10.1093/nar/gkv1003

Carten JD, Bradford MK, Farber SA. 2011. Visualizing digestive organ morphology and function using differential fatty acid metabolism in live zebrafish. Dev Biol. doi:10.1016/j.ydbio.2011.09.010

Chinnery PF, Elliott HR, Hudson G, Samuels DC, Relton CL. 2012. Epigenetics, epidemiology and mitochondrial DNA diseases. Int J Epidemiol 41:177–187. doi:10.1093/ije/dyr232

Chujo T, Ohira T, Sakaguchi Y, Goshima N, Nomura N, Nagao A, Suzuki T. 2012. LRPPRC/SLIRP suppresses PNPase-mediated mRNA decay and promotes polyadenylation in human mitochondria. Nucleic Acids Res 40:8033–8047. doi:10.1093/nar/gks506

Clark KJ, Balciunas D, Pogoda H-M, Ding Y, Westcot SE, Bedell VM, Greenwood TM, Urban MD, Skuster KJ, Petzold AM, Ni J, Nielsen AL, Patowary A, Scaria V, Sivasubbu S, Xu X, Hammerschmidt M, Ekker SC. 2011. In vivo protein trapping produces a functional expression codex of the vertebrate proteome. Nat Methods 8:506–515. doi:10.1038/nmeth.1606

Cui J, Wang L, Ren X, Zhang Y, Zhang H. 2019. LRPPRC: A Multifunctional Protein Involved in Energy Metabolism and Human Disease. Front Physiol. doi:10.3389/fphys.2019.00595

Cuillerier A, Honarmand S, Cadete VJJ, Ruiz M, Forest A, Deschenes S, Beauchamp C, Charron G, Rioux JD, Des Rosiers C, Shoubridge EA, Burelle Y. 2017. Loss of hepatic LRPPRC alters mitochondrial bioenergetics, regulation of permeability transition and trans-membrane ROS diffusion. Hum Mol Genet 26:3186–3201. doi:10.1093/hmg/ddx202

Debray F-G, Morin C, Janvier A, Villeneuve J, Maranda B, Laframboise R, Lacroix J, Decarie J-C, Robitaille Y, Lambert M, Robinson BH, Mitchell GA. 2011. LRPPRC mutations cause a phenotypically distinct form of Leigh syndrome with cytochrome c oxidase deficiency. J Med Genet 48:183–189. doi:10.1136/jmg.2010.081976

Dominguez-Perez M, Simoni-Nieves A, Rosales P, Nuno-Lambarri N, Rosas-Lemus M, Souza V, Miranda RU, Bucio L, Uribe Carvajal S, Marquardt JU, Seo D, Gomez-Quiroz LE, Gutierrez-Ruiz MC. 2019. Cholesterol burden in the liver induces mitochondrial dynamic changes and resistance to apoptosis. J Cell Physiol 234:7213–7223. doi:10.1002/jcp.27474

Flinn L, Mortiboys H, Volkmann K, Kster RW, Ingham PW, Bandmann O. 2009. Complex i deficiency and dopaminergic neuronal cell loss in parkin-deficient zebrafish (Danio rerio). Brain. doi:10.1093/brain/awp108

Glover LE, Newton K, Krishnan G, Bronson R, Boyle A, Krivickas LS, Brown RHJ. 2010. Dysferlin overexpression in skeletal muscle produces a progressive myopathy. Ann Neurol 67:384–393. doi:10.1002/ana.21926

Gohil VM, Nilsson R, Belcher-Timme CA, Luo B, Root DE, Mootha VK. 2010. Mitochondrial and nuclear genomic responses to loss of LRPPRC expression. J Biol Chem 285:13742–13747. doi:10.1074/jbc.M109.098400

Gorman GS, Chinnery PF, DiMauro S, Hirano M, Koga Y, McFarland R, Suomalainen A, Thorburn DR, Zeviani M, Turnbull DM. 2016. Mitochondrial diseases. Nat Rev Dis Prim 2:16080. doi:10.1038/nrdp.2016.80

Green DR. 1998. Apoptotic pathways: the roads to ruin. Cell 94:695–698. doi:10.1016/s0092-8674(00)81728-6

Henderson JR, Macalma T, Brown D, Richardson JA, Olson EN, Beckerle MC. 1999. The LIM protein, CRP1, is a smooth muscle marker. Dev Dyn 214:229–238. doi:10.1002/(SICI)1097-0177(199903)214:3<229::AID-AJA6>3.0.CO;2-S

Ichino N, Serres M, Urban R, Urban M, Schaefbauer K, Greif L, Varshney GK, Skuster KJ, McNulty M, Daby C, Wang Y, Liao H, El-Rass S, Ding Y, Liu W, Schimmenti LA, Sivasubbu S, Balciunas D, Hammerschmidt M, Farber SA, Wen X-Y, Xu X, McGrail M, Essner JJ, Burgess S, Clark KJ, Ekker SC. 2019. The Vertebrate Codex Gene Breaking Protein Trap Library For Genomic Discovery and Disease Modeling Applications. bioRxiv 630236. doi:10.1101/630236

Iida R, Yasuda T, Tsubota E, Takatsuka H, Matsuki T, Kishi K. 2006. Human Mpv17-like protein is localized in peroxisomes and regulates expression of antioxidant enzymes. Biochem Biophys Res Commun 344:948–954. doi:10.1016/j.bbrc.2006.04.008

Kim MJ, Kang KH, Kim CH, Choi SY. 2008. Real-time imaging of mitochondria in transgenic zebrafish expressing mitochondrially targeted GFP. Biotechniques. doi:10.2144/000112909

Kim S-H, Scott SA, Bennett MJ, Carson RP, Fessel J, Brown HA, Ess KC. 2013. Multi-organ abnormalities and mTORC1 activation in zebrafish model of multiple acyl-CoA dehydrogenase deficiency. PLoS Genet 9:e1003563. doi:10.1371/journal.pgen.1003563

Kohler F, Muller-Rischart AK, Conradt B, Rolland SG. 2015. The loss of LRPPRC function induces the mitochondrial unfolded protein response. Aging (Albany NY) 7:701–717. doi:10.18632/aging.100812

Lambert CJ, Freshner BC, Chung A, Stevenson TJ, Bowles DM, Samuel R, Gale BK, Bonkowsky JL. 2018. An automated system for rapid cellular extraction from live zebrafish embryos and larvae: Development and application to genotyping. PLoS One 13:e0193180. doi:10.1371/journal.pone.0193180

Lieschke GJ, Currie PD. 2007. Animal models of human disease: Zebrafish swim into view. Nat Rev Genet. doi:10.1038/nrg2091

Liu L, McKeehan WL. 2002. Sequence analysis of LRPPRC and its SEC1 domain interaction partners suggests roles in cytoskeletal organization, vesicular trafficking, nucleocytosolic shuttling, and chromosome activity. Genomics. doi:10.1006/geno.2001.6679

Love MI, Huber W, Anders S. 2014. Moderated estimation of fold change and dispersion for RNA-seq data with DESeq2. Genome Biol 15:550. doi:10.1186/s13059-014-0550-8

Manna S. 2015. An overview of pentatricopeptide repeat proteins and their applications. Biochimie. doi:10.1016/j.biochi.2015.04.004

McFarland R, Taylor RW, Turnbull DM. 2010. A neurological perspective on mitochondrial disease. Lancet Neurol 9:829–840. doi:10.1016/S1474-4422(10)70116-2

Mi H, Muruganujan A, Huang X, Ebert D, Mills C, Guo X, Thomas PD. 2019. Protocol Update for large-scale genome and gene function analysis with the PANTHER classification system (v.14.0). Nat Protoc 14:703–721. doi:10.1038/s41596-019-0128-8

Mili S, Piñol-Roma S. 2003. LRP130, a Pentatricopeptide Motif Protein with a Noncanonical RNA-Binding Domain, Is Bound In Vivo to Mitochondrial and Nuclear RNAs. Mol Cell Biol. doi:10.1128/mcb.23.14.4972-4982.2003

Mootha VK, Lepage P, Miller K, Bunkenborg J, Reich M, Hjerrild M, Delmonte T, Villeneuve A, Sladek R, Xu F, Mitchell GA, Morin C, Mann M, Hudson TJ, Robinson B, Rioux JD, Lander ES. 2003. Identification of a gene causing human cytochrome c oxidase deficiency by integrative genomics. Proc Natl Acad Sci U S A 100:605–610. doi:10.1073/pnas.242716699

Moraes CT, Ricci E, Bonilla E, DiMauro S, Schon EA. 1992. The mitochondrial tRNA(Leu(UUR)) mutation in mitochondrial encephalomyopathy, lactic acidosis, and strokelike episodes (MELAS): genetic, biochemical, and morphological correlations in skeletal muscle. Am J Hum Genet 50:934–949.

Morin C, Mitchell G, Larochelle J, Lambert M, Ogier H, Robinson BH, De Braekeleer M. 1993. Clinical, metabolic, and genetic aspects of cytochrome C oxidase deficiency in Saguenay-Lac-Saint-Jean. Am J Hum Genet 53:488–496.

Morino K, Petersen KF, Shulman GI. 2006. Molecular mechanisms of insulin resistance in humans and their potential links with mitochondrial dysfunction. Diabetes 55 Suppl 2:S9–S15. doi:10.2337/db06-S002

Nene A, Chen C-H, Disatnik M-H, Cruz L, Mochly-Rosen D. 2017. Aldehyde dehydrogenase 2 activation and coevolution of its epsilonPKC-mediated phosphorylation sites. J Biomed Sci 24:3. doi:10.1186/s12929-016-0312-x

Newsholme P, Gaudel C, Krause M. 2012. Mitochondria and diabetes. An intriguing pathogenetic role. Adv Exp Med Biol 942:235–247. doi:10.1007/978-94-007-2869-1_10

Olahova M, Hardy SA, Hall J, Yarham JW, Haack TB, Wilson WC, Alston CL, He L, Aznauryan E, Brown RM, Brown GK, Morris AAM, Mundy H, Broomfield A, Barbosa IA, Simpson MA, Deshpande C, Moeslinger D, Koch J, Stettner GM, Bonnen PE, Prokisch H, Lightowlers RN, McFarland R, Chrzanowska-Lightowlers ZMA, Taylor RW. 2015. LRPPRC mutations cause early-onset multisystem mitochondrial disease outside of the French-Canadian population. Brain 138:3503–3519. doi:10.1093/brain/awv291

Plucinska G, Paquet D, Hruscha A, Godinho L, Haass C, Schmid B, Misgeld T. 2012. In vivo imaging of disease-related mitochondrial dynamics in a vertebrate model system. J Neurosci. doi:10.1523/JNEUROSCI.1327-12.2012

Quinlivan VH, Wilson MH, Ruzicka J, Farber SA. 2017. An HPLC-CAD/fluorescence lipidomics platform using fluorescent fatty acids as metabolic tracers. J Lipid Res 58:1008–1020. doi:10.1194/jlr.D072918

Rahman S, Blok RB, Dahl HH, Danks DM, Kirby DM, Chow CW, Christodoulou J, Thorburn DR. 1996. Leigh syndrome: clinical features and biochemical and DNA abnormalities. Ann Neurol 39:343–351. doi:10.1002/ana.410390311

Rizzuto R, Marchi S, Bonora M, Aguiari P, Bononi A, De Stefani D, Giorgi C, Leo S, Rimessi A, Siviero R, Zecchini E, Pinton P. 2009. Ca(2+) transfer from the ER to mitochondria: when, how and why. Biochim Biophys Acta 1787:1342–1351. doi:10.1016/j.bbabio.2009.03.015

Ruiz M, Cuillerier A, Daneault C, Deschenes S, Frayne IR, Bouchard B, Forest A, Legault JT, Vaz FM, Rioux JD, Burelle Y, Des Rosiers C. 2019. Lipidomics unveils lipid dyshomeostasis and low circulating plasmalogens as biomarkers in a monogenic mitochondrial disorder. JCI insight 4. doi:10.1172/jci.insight.123231

Ruzzenente B, Metodiev MD, Wredenberg A, Bratic A, Park CB, Camara Y, Milenkovic D, Zickermann V, Wibom R, Hultenby K, Erdjument-Bromage H, Tempst P, Brandt U, Stewart JB, Gustafsson CM, Larsson N-G. 2012. LRPPRC is necessary for polyadenylation and coordination of translation of mitochondrial mRNAs. EMBO J 31:443–456. doi:10.1038/emboj.2011.392

Sabharwal A, Campbell JM, WareJoncas Z, Wishman M, Ata H, Liu W, Ichino N, Bergren JD, Urban MD, Urban R, Poshusta TL, Ding Y, Xu X, Clark KJ, Ekker SC. 2019a. A primer genetic toolkit for exploring mitochondrial biology and disease using zebrafish. bioRxiv. doi:10.1101/542084

Sabharwal A, Sharma D, Vellarikkal SK, Jayarajan R, Verma A, Senthivel V, Scaria V, Sivasubbu S. 2019b. Organellar transcriptome sequencing reveals mitochondrial localization of nuclear encoded transcripts. Mitochondrion 46:59–68. doi:10.1016/j.mito.2018.02.007

Sasarman F, Nishimura T, Antonicka H, Weraarpachai W, Shoubridge EA. 2015. Tissue-specific responses to the LRPPRC founder mutation in French Canadian Leigh Syndrome. Hum Mol Genet 24:480–491. doi:10.1093/hmg/ddu468

Siira SJ, Spahr H, Shearwood A-MJ, Ruzzenente B, Larsson N-G, Rackham O, Filipovska A. 2017. LRPPRC-mediated folding of the mitochondrial transcriptome. Nat Commun 8:1532. doi:10.1038/s41467-017-01221-z

Simoes ICM, Fontes A, Pinton P, Zischka H, Wieckowski MR. 2018. Mitochondria in non-alcoholic fatty liver disease. Int J Biochem Cell Biol 95:93–99. doi:10.1016/j.biocel.2017.12.019

Smith LL, Beggs AH, Gupta VA. 2013. Analysis of skeletal muscle defects in larval zebrafish by birefringence and touch-evoke escape response assays. J Vis Exp. doi:10.3791/50925

Soneson C, Love MI, Robinson MD. 2015. Differential analyses for RNA-seq: transcript-level estimates improve gene-level inferences. F1000Research 4:1521. doi:10.12688/f1000research.7563.2

Song Y, Selak MA, Watson CT, Coutts C, Scherer PC, Panzer JA, Gibbs S, Scott MO, Willer G, Gregg RG, Ali DW, Bennett MJ, Balice-Gordon RJ. 2009. Mechanisms underlying metabolic and neural defects in zebrafish and human multiple Acyl-CoA dehydrogenase deficiency (MADD). PLoS One. doi:10.1371/journal.pone.0008329

Spinelli JB, Haigis MC. 2018. The multifaceted contributions of mitochondria to cellular metabolism. Nat Cell Biol 20:745–754. doi:10.1038/s41556-018-0124-1

Steele SL, Prykhozhij S V., Berman JN. 2014. Zebrafish as a model system for mitochondrial biology and diseases. Transl Res. doi:10.1016/j.trsl.2013.08.008

Sterky FH, Ruzzenente B, Gustafsson CM, Samuelsson T, Larsson N-G. 2010. LRPPRC is a mitochondrial matrix protein that is conserved in metazoans. Biochem Biophys Res Commun 398:759–764. doi:10.1016/j.bbrc.2010.07.019

Stockel D, Kehl T, Trampert P, Schneider L, Backes C, Ludwig N, Gerasch A, Kaufmann M, Gessler M, Graf N, Meese E, Keller A, Lenhof H-P. 2016. Multi-omics enrichment analysis using the GeneTrail2 web service. Bioinformatics 32:1502–1508. doi:10.1093/bioinformatics/btv770

Sun S, Li H, Chen J, Qian Q. 2017. Lactic Acid: No Longer an Inert and End-Product of Glycolysis. Physiology (Bethesda) 32:453–463. doi:10.1152/physiol.00016.2017

Tiku V, Tan M-W, Dikic I. 2020. Mitochondrial Functions in Infection and Immunity. Trends Cell Biol 30:263–275. doi:10.1016/j.tcb.2020.01.006

Urrutia PJ, Mena NP, Nunez MT. 2014. The interplay between iron accumulation, mitochondrial dysfunction, and inflammation during the execution step of neurodegenerative disorders. Front Pharmacol 5:38. doi:10.3389/fphar.2014.00038

Vafai SB, Mootha VK. 2012. Mitochondrial disorders as windows into an ancient organelle. Nature 491:374–383. doi:10.1038/nature11707

Valero T. 2014. Mitochondrial biogenesis: pharmacological approaches. Curr Pharm Des. doi:10.2174/138161282035140911142118

Wallace DC. 2012. Mitochondria and cancer. Nat Rev Cancer 12:685–698. doi:10.1038/nrc3365

Wilkinson RN, Elworthy S, Ingham PW, van Eeden FJM. 2017. Fin clipping and genotyping embryonic zebrafish at 3 days post-fertilization. Biotechniques. doi:10.2144/000114509

Wilson MH, Rajan S, Danoff A, White RJ, Hensley MR, Quinlivan VH, Thierer JH, Busch-Nentwich EM, Hussain MM, Farber SA. 2019. A missense mutation dissociates triglyceride and phospholipid transfer activities in zebrafish and human microsomal triglyceride transfer protein. bioRxiv 701813. doi:10.1101/701813

Xu F, Addis JBL, Cameron JM, Robinson BH. 2012. LRPPRC mutation suppresses cytochrome oxidase activity by altering mitochondrial RNA transcript stability in a mouse model. Biochem J 441:275–283. doi:10.1042/BJ20110985

Zeituni EM, Wilson MH, Zheng X, Iglesias PA, Sepanski MA, Siddiqi MA, Anderson JL, Zheng Y, Farber SA. 2016. Endoplasmic reticulum lipid flux influences enterocyte nuclear morphology and lipid-dependent transcriptional responses. J Biol Chem. doi:10.1074/jbc.M116.749358

